# Geometrical preference of anchoring sites in the unicellular organism *Stentor coeruleus*

**DOI:** 10.1101/2025.07.20.664703

**Authors:** Syun Echigoya, Takuya Ohmura, Katsuhiko Sato, Toshiyuki Nakagaki, Yukinori Nishigami

## Abstract

Organisms often inhabit environments comprising complex structures across various scales. Animals rely on visual information from surrounding geometrical structures for navigation. Even at the microscale, various microsediments form complex structures in microbial habitats. The movement of microorganisms is passively affected by collisions and hydrodynamic interactions with surrounding structures. However, the influence of microenvironmental geometry on behavioral changes of unicellular organisms that lack visual perception remains unclear. Here, we developed geometrically structured chambers to investigate anchoring site preferences in the swimming ciliate *Stentor coeruleus*. Our experiments revealed that *S. coeruleus* preferentially anchored in narrow regions characterized by specific geometrical features, including corner angle, depth, and curvature at the corner end. Before anchoring, free-swimming *S. coeruleus* changed its behavior to move along the boundary wall of the chambers, accompanied by Ca^2+^-induced asymmetrical body deformation. To further investigate how *S. coeruleus* moves along the wall continuously, we conducted a hydrodynamic simulation and revealed that the asymmetric morphology causes asymmetric propulsive forces, explaining wall-following behavior through physical interactions with a wall. Thus, morphological change near a wall causes wall-following behavior, facilitating the identification of these narrow anchoring sites. Our findings indicate that environmental geometry drives behavioral transitions in *S. coeruleus* through simple biophysical processes, enabling spatial selection without visual cues. Overall, these results suggest that microgeometry plays a key role in shaping ecological niches for unicellular microorganisms.

**Significance Statement:** Animals use various natural structures as landmarks for navigation. In microorganism habitats, microsediments also form geometrically complex environments. Is there a relationship between the geometrical features of structures and the behavior in unicellular organisms lacking visual cues? Here we report that the free-swimming unicellular organism *Stentor coeruleus* selects the anchoring sites based on the surrounding shapes. Further observations and numerical simulations reveal that an asymmetric morphological change causes a temporary switch from ballistic to wall-following exploration, driven by surrounding structures. These results indicate that one simple behavioral response underlies the preference of anchoring sites with specific geometrical features in non-neural unicellular organisms. The findings shed light on the role of microenvironmental geometry in forming ecological niches for microorganisms.

## Introduction

Navigation is a fundamental survival strategy for heterogeneous environments. Animals, such as insects and mammals, use global information from the sun, light polarization, magnetic fields, and gravity for navigation (1–5). Moreover, ants, bees, and mantis shrimp use various natural structures in their habitats as panoramic views or landmarks for navigation (6–9). In a recent study, environmental geometry obtained through visual input was encoded by a single cell in the dorsal subiculum in mice (10). Thus, various behavioral experiments and electrophysiological analyses have suggested that ubiquitous obstacles and their geometrical shapes play important roles in animal navigation (11).

Spatial exploration in unicellular organisms has been studied based on the taxis of primitive navigation systems that react to light, magnetic fields, electric fields, gravity, and chemical substances (12–14). Spatiotemporal changes in external stimulus cause Ca^2+^ influx into cilia through channels (15, 16). These bioelectrical signals modulate ciliary beating patterns and motility (17–21), leading to tactic behaviors (22). Active regulation of ciliary beating in algae and shape changes in ciliates exhibit adaptive responses originating from interactions with hydrodynamic constraints (23, 24).

In microorganism habitats, many microsediments form geometrically complex environments. In the last decade, the physical effect of geometrical structures on the behavior of microorganisms has been investigated through mechanical collision or hydrodynamic interaction (25–34). These studies examined passive scattering events arising from interactions with planar and curved boundaries, fixed-angle corners, and avoidance responses in crowded pillar arrays. However, natural environments present far more complex geometries in which multiple, competing geometric alternatives coexist within the same habitat. How a unicellular organism behaves under such increased geometrical complexity remains poorly understood. Consequently, it is unclear to what extent microorganisms can sense and respond to geometrical features of their environment.

The swimming ciliate, *Stentor coeruleus*, exhibits complex behavior that switches between free-swimming and anchoring to a substrate (Fig. 1A). In the swimming state, *S. coeruleus* generates propulsive force primarily through hair-like organelles, termed membranellar band (MB), located around the anterior. The cell explores a space with its trajectory varying in response to light and chemical cues (35, 36). During swimming, *S. coeruleus* gradually elongates into a trumpet shape and adheres to a substrate via an anchoring organ (holdfast) at its posterior end (32, 37). Anchored *S. coeruleus* also generates external vortical flows through MB, forming an oral apparatus for capturing bacteria and small ciliates as food (38). At the same time, attachment may increase the risk of predation. Thus, selecting anchoring sites in heterogeneous environments may represent an essential behavior in *S. coeruleus*.

**Fig. 1.**
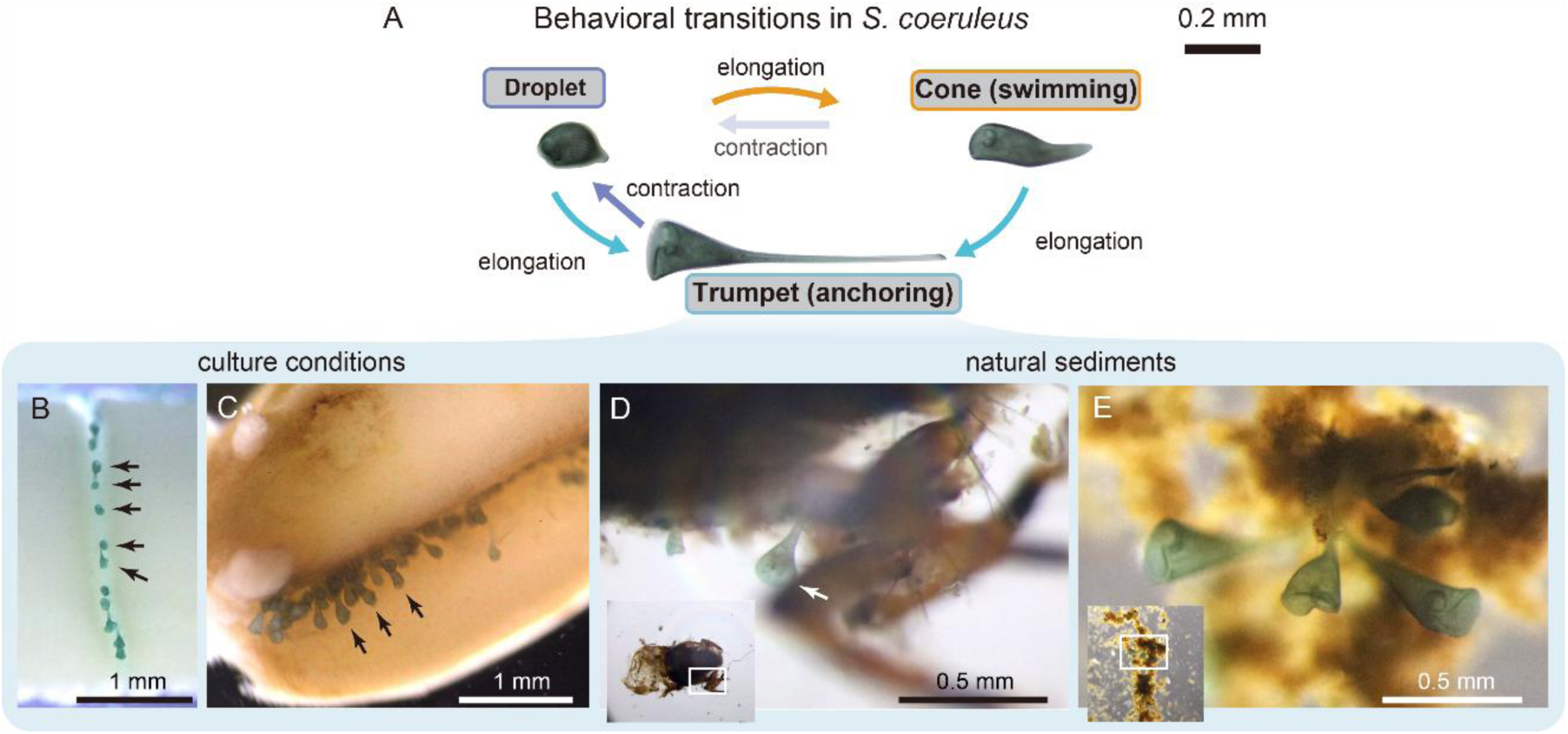
Anchoring sites of *Stentor coeruleus* in natural and laboratory conditions. **(A)** Three types of behavioral modes accompanying shape change in the ciliate *S. coeruleus*. *S. coeruleus* alternates between free swimming and attachment to a substrate. A brief mechanical stimulus induces rapid contraction into a droplet-like shape as an avoidance response. Modified from Echigoya et al (32). **(B, C)** Anchoring sites in culture conditions, such as the channel on a rice grain (B) and an oat grain (C). Each arrow indicates one anchoring *S. coeruleus*. **(D**, **E)** Anchoring sites in natural sediments. *S. coeruleus* anchored to a narrow area in the larval exuvia of a caddisfly (Trichoptera) under laboratory conditions (D). Trumpet-shaped anchoring *S. coeruleus* were frequently observed on complex sediments upon sample collection from natural environment (E). The rectangles in the insets represent the focused regions.

Under culture conditions, we often found *S. coeruleus* anchoring to rice or oat grains added to promote bacterial growth as a nutrient source (Fig. 1B, C). Under laboratory conditions, *S. coeruleus* also anchored in narrow sites in natural sediments, such as the exuvia of an aquatic insect (Fig. 1D). When sampling from a pond, anchoring *S. coeruleus* were often found on a geometrically complex sediment (Fig. 1E). *S. coeruleus* tended to anchor within narrow areas and not across the entire surface of the sediments (Fig. 1B–E). The environmental complexity in both culture and natural conditions made it difficult to identify the factors influencing the selection of anchoring sites.

Herein, we developed three types of geometrically structured chambers using a sub-millimeter-scale fabrication technique to investigate the anchoring behavior of *S. coeruleus*, extending our previous study (32). *S. coeruleus* exhibited a geometrical preference of anchoring sites based on the corner angle, depth, and curvature at the corner ends formed by a structure and chamber walls. This geometrical preference exhibited a correlation with body length in the swimming state. Before anchoring in the geometrical structural chambers, *S. coeruleus* displayed a behavioral change from exploratory to wall-following behavior, which was physically explained by asymmetric propulsion accompanying cell deformation regulated by intracellular Ca^2+^. This study addresses how environmental geometry influences exploratory and anchoring behavior, shedding light on how complex environments guide site selection of unicellular organisms in narrow areas.

## Results

### Geometrical preferences of anchoring sites in geometrically complex environments

We developed quasi-2D geometrically structured chambers to investigate the geometrical features of the anchoring sites of *S. coeruleus*. We observed the behavior of a single *S. coeruleus* for approximately 40 min under dim red light to prevent phototactic responses in these chambers (Fig. 2A). *S. coeruleus* freely swam with a spiral trajectory exploring the chambers. The swimming speed temporarily decreased around the walls and corners owing to collisions caused by the geometrical complexity of the chambers (Fig. 2B). To analyze the overview of the behaviors of *S. coeruleus*, we quantified the swimming speed and cell length. Most cells exhibited a length of 0.4 mm and swam steadily at a speed of 0.5–1.0 mm/s. Subsequently, the swimming cells switched to the anchoring state accompanied by elongation and deceleration (Fig. 2C and Movie S1). Some cells repeatedly switched their states between swimming and anchoring, leading to behavioral complexity. We measured the locations of the cells and anchoring rate based on the number of anchoring events, focusing on the three geometrical features: corner angle, depth, and curvature.

**Fig. 2.**
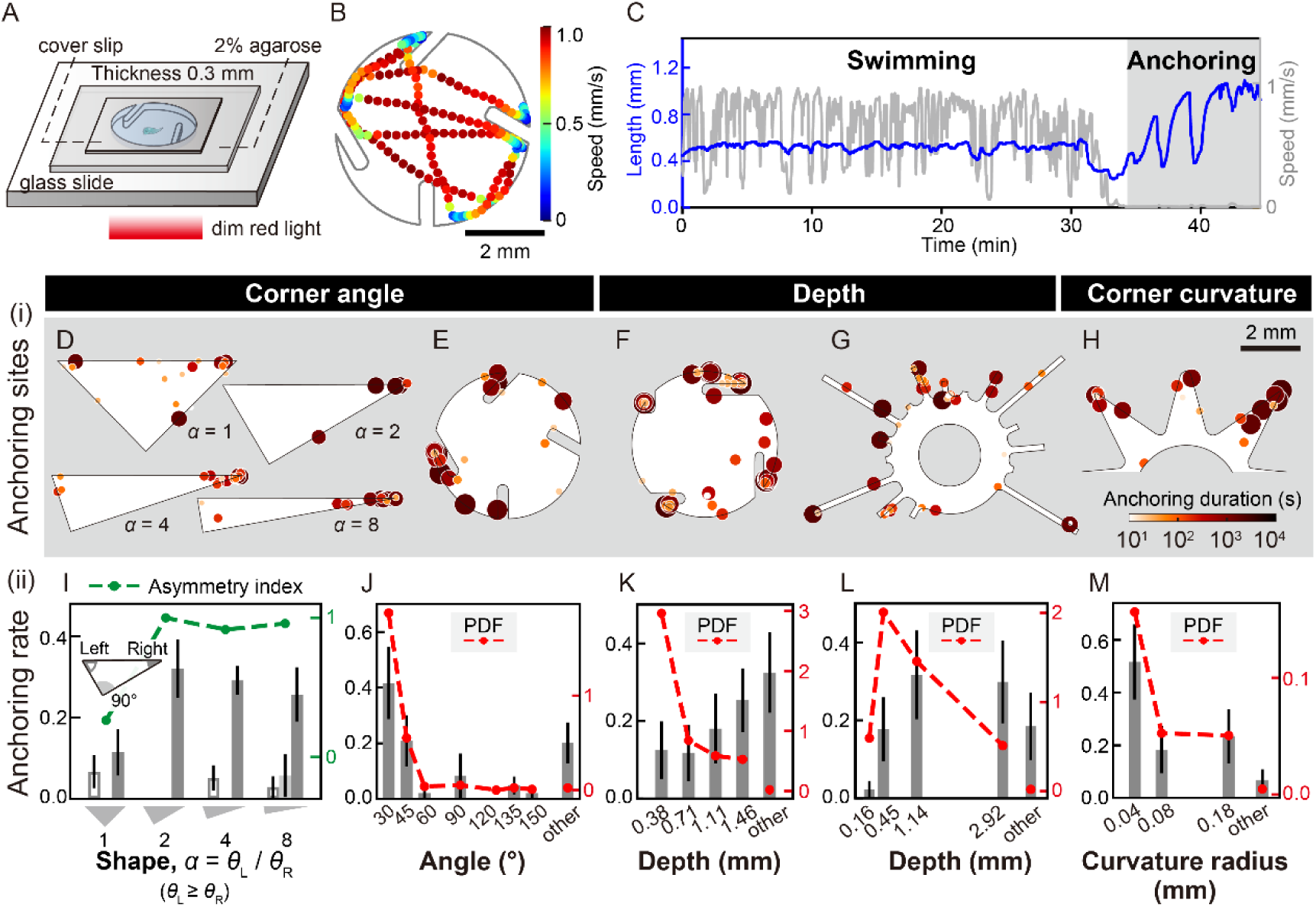
Geometrical preferences of anchoring sites in geometrically complex environments in *Stentor coeruleus*. **(A)** Experimental setup for behavioral observation in geometrical structural chambers. The behavior of *S. coeruleus* was observed in the chamber with a microstructure made of agarose gel and sealed with coverslip. Observation were conducted under weak red light to avoid phototactic responses in *S. coeruleus*. **(B)** Swimming trajectory of *S. coeruleus* in a steady state in a geometrically structured chamber. Cell positions recorded every 0.25 s are plotted as points. Colors represent swimming speed. **(C)** Overview of the behaviors focusing on the time series for swimming speed (gray line) and cell length (blue line), respectively. Swimming speed gradually decreased to zero as the cell transitioned from swimming to anchoring. This behavioral change was accompanied by a gradual elongation of the cell. **(D–H)** Anchoring sites in several geometrically structured chambers. Each dot corresponds to an anchoring site. Marker size visually corresponds to color, which represents the duration of an anchoring event. **(I–M)** Quantitative analysis of anchoring events in each specific area with different geometrical features within a chamber type. The green dashed line represents the asymmetry index calculated from anchoring rate divided by anchoring area (I). Red dashed lines indicate the probability density functions (PDFs), calculated as anchoring rate divided by anchoring area (J–M). The definitions of specific areas are included in SI Appendix, Fig. S1A–E. *n* = 7–9 cells in each chamber (D), *n* = 12 cells (E), *n* = 20 cells (F), *n* = 18 cells (G), and *n* = 15 cells (H). See also SI Appendix, Figs. S1–S3 and Movie S1.

First, in a symmetrical right-triangle chamber, *S. coeruleus* anchored more frequently at 45° corners than at a 90° corner, and the number of anchoring events in each 45° corner was not significantly different (Fig. 2D, I and SI Appendix, Fig. S2B, binomial test, *p* = 0.36). In an asymmetrical right-triangle chamber, *S. coeruleus* anchored to a corner with a small angle (Fig. 2D). Next, we designed an environment containing corner angles ranging from 30° to 150° by introducing protruding rod-like structures along the outer edge of the chambers. Subsequently, the cells showed a tendency to anchor at corners with small angles (Fig. 2E). Although 27% of anchoring events (6/22) were observed outside the corners, probability densities calculated as anchoring rates per area were relatively low compared with those at corners with acute angles (Fig. 2J). Thus, corners with angles less than 45° showed a significant increase in anchoring rate and total anchoring duration (Fig. 2J and SI Appendix, Fig. S2).

Second, we developed a geometrically structured chamber featuring four types of corners, each having a depth and protruding rod-like structures with different lengths at a fixed angle. In the chamber, *S. coeruleus* tended to anchor to corners with a greater depth (Fig. 2F and SI Appendix, Fig. S2). Although anchoring rate and total anchoring duration exhibited a positive correlation with the corner depth, several cells anchored at the corner formed by the short protrusion (depth: ∼0.4 mm) (Fig. 2K). The probability density of anchoring was higher at this corner than at other corners (Fig. 2K). In channels with different lengths, *S. coeruleus* anchored at channels deeper than 0.4 mm (Fig. 2G, L). However, the cells did not frequently anchor to the channels with a depth more than twice their cell length; the highest probability density of anchoring was observed at a depth of 0.45 mm (Fig. 2L). Notably, while the cells spent 75% of their total swimming time outside the channels, 85% of the anchoring events (33/39) were observed inside one of the channels (SI Appendix, Fig. S2).

Third, we arranged three types of corners with different two-dimensional curvatures in a star-shaped periodic layout at three locations. Fig. 2H represents the superimposed anchoring sites and durations at each corner type, showing a tendency to anchor near corners with a smaller curvature radius (<0.1 mm), which corresponds to the width of *S. coeruleus* (Fig. 2M). In addition, we investigated whether a hierarchy of geometrical features influenced the selection of anchoring sites by observing the anchoring behavior of these cells at corners with different angles and curvatures (SI Appendix, Fig. S2). Most cells anchored at sharper corners with a smaller angle. However, the secondary preferences in the selection of anchoring sites were unclear, suggesting no hierarchical prioritization of the anchoring sites.

Collectively, the aforementioned findings demonstrate that *S. coeruleus* selects anchoring sites based on the surrounding geometrical features and that the geometrical preferences scale with its cell size in the swimming state.

### Behavioral process of anchoring in geometrically complex environments

To understand the anchoring behavior in the geometrically structured chambers, we focused on the behavioral transition from steady swimming to the trumpet-shaped anchored state and compared this transition between a simple circular chamber and geometrically structured chambers. We found two types of anchoring processes depending on the structural complexity of the environment (Fig. 3A). In the simple circular chamber, the cell monotonically elongated from a conical swimming shape to a trumpet shape during anchoring, consistent with previous reports (32, 37). In comparison, in the geometrically structured chambers, swimming *S. coeruleus* exhibited a transient contraction over a few seconds before elongation. This contraction in the geometrically structured chambers reduced the cell length to approximately 70% of the original, representing a significant decrease compared with the cell length in the simple circular chamber 30 s after each transition (Fig. 3B). The characteristic time of the transition from contraction to elongation was not observed owing to variability in the transition time among individual cells. Even in the geometrically structured chambers, some cells monotonically elongated in the anchoring transitions (Fig. 3A, B). The cellular length in the final state, measured as the maximum value in each transition, showed no significant difference in the anchoring processes across the three conditions (Fig. 3B).

**Fig. 3.**
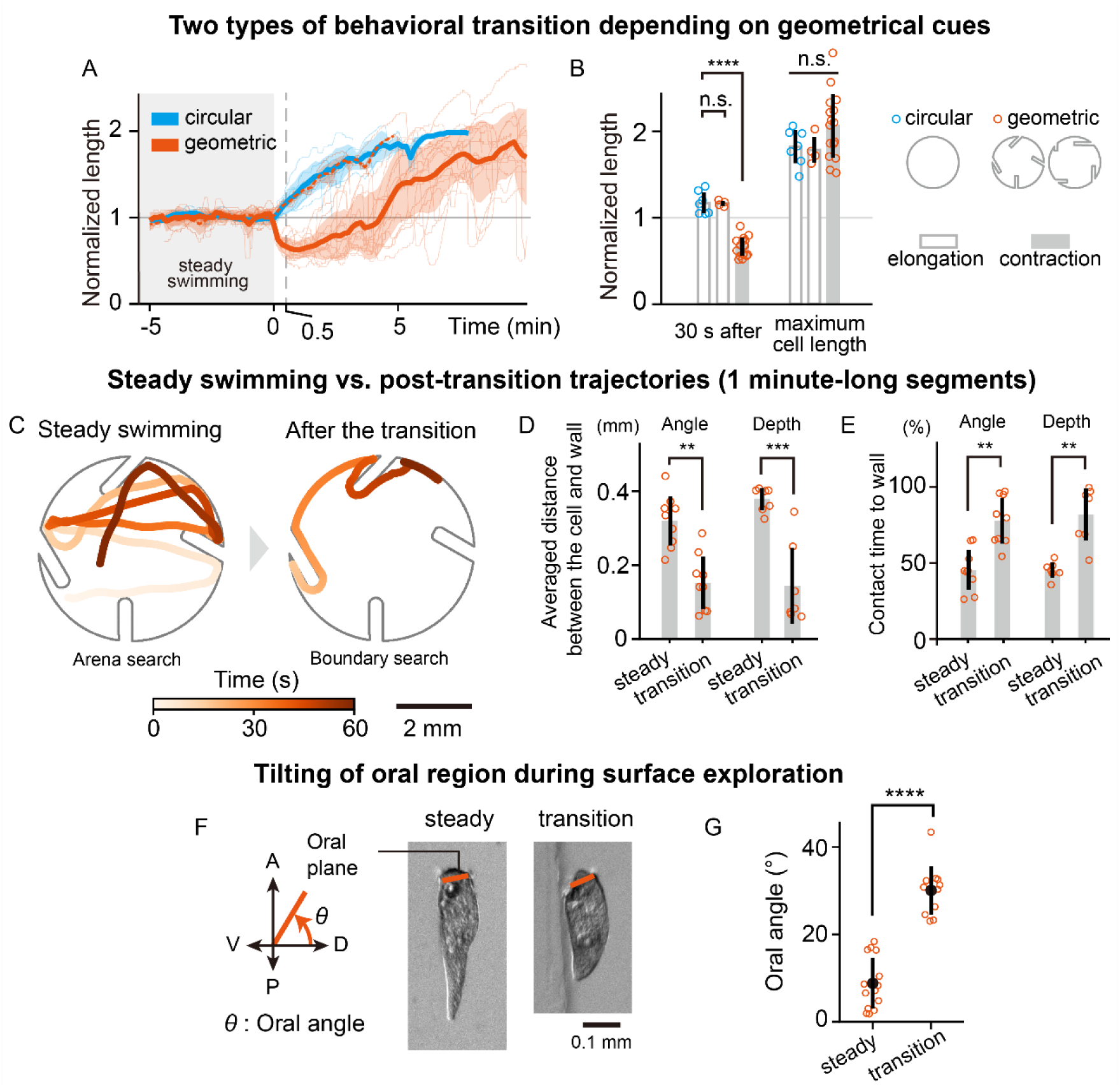
Behavioral process of anchoring influenced by asymmetric deformation of body shape in geometrically structured chambers. **(A)** Time series of cellular length in two behavioral transitions in the chamber with/without a geometrical structure. Cellular lengths were standardized by each individual mean pre-transition length, with the onset of transition aligned to *t* = 0. The thick solid lines and the thin colored areas correspond to the mean length and the standard deviation (SD) of the length, respectively. *n* = 16, contraction in the geometrically structured chambers; *n* = 4, elongation in the geometrically structured chambers; and *n* = 7 elongation in the simple circular chamber. **(B)** Comparison of the normalized lengths in two transition types between the simple circular and geometrically structured chambers. The lengths 30 s after the transition were significantly different between the two behavioral processes (Welch’s *t*-test; *p* < 10^−5^; *t* = 8.97; one side; *n* = 7; *n* = 16). The lengths 30 s after the monotonic elongation process were not significantly different between the simple circular and geometrically structured chambers (Welch’s *t*-test; *p* = 0.93; *t* = 0.095; *n* = 7; *n* = 4). There was no significant difference in the maximum normalized cell lengths after the transition (one-way ANOVA; *p* = 0.15; *F* (2, 24) = 2.05). **(C)** Typical swimming trajectories over 1 min before and after the transition in the geometrically structured chamber. Color intensity represents time progression. Cells transitioned from broad exploration to surface exploration behavior. **(D)** Quantification of surface exploration calculated as the mean distance between the wall and cell center in two geometrically structured chambers (paired *t*-test, angle; *p* = 0.0024, *t* = −3.87, one side, *n* = 9 cells; depth, *p* = 0.00068, *t* = −5.62, one side, *n* = 7 cells). **(E)** Quantification of surface exploration calculated as the averaged ratio of wall-contact time to total time during the behavioral window of up to 1 minute (paired *t*-test, angle chamber; *p* = 0.0044, *t* = 3.45, one-side, 9 cells; depth chamber, *p* = 0.0014, *t* = 4.87, one side, *n* = 7 cells). **(F, G)** Measurement of the oral plane of the cell. The tilted oral angle is defined as the angle between the anterior vector and the adjusted vector of the oral plane. The dorsal-to-anterior direction is a positive angle (F). Statistical analysis showed a significant difference in the tilted angle between the steady state and the transition state (G) (Welch’s *t*-test, *p* < 10^−8^, *t* = −8.92, one side, *n* = 14 cells in the steady state, *n* = 11 cells in the transition state). All error bars represent SDs. See also SI Appendix, Figs S4, S6, Movies S2 and S3.

To further understand the anchoring transition with temporal contraction, we examined the differences in swimming trajectories between the steady swimming state and the post-transition state within a behavioral window of up to 1 minute. In the steady swimming state, the cells explored the whole geometrically structured chambers, with the chamber wall affecting the swimming directions (Movie S2). A helical swimming trajectory was observed, accompanied by rotation around the long axis of the cell. After transitioning, cells adopted wall-following swimming behaviors, moving along even rod-like structures without rotation around their swimming direction (Fig. 3C, SI Appendix, Fig. S4, and Movie S3). Compared with steady swimming cells, post-transition cells moved closer to the chamber wall, as indicated by the average distance between the cell center and the wall in two geometrically structured chambers (Fig. 3D). Furthermore, the ratio of wall-contact time to total time during the behavioral window following the transition was also significantly high (Fig. 3E). Thus, during the anchoring process in the geometrically structured chambers, *S. coeruleus* exhibits a wall-following behavior without helical swimming after transient contraction, which precedes cell elongation.

### Effect of intracellular Ca^2+^ concentration on the anchoring process accompanying cell shape changes

In the class Heterotrichea, including *Stentor*, cell deformation is regulated by intracellular Ca^2+^ concentration (39, 40). Therefore, to understand the biological mechanism of cell deformation during the anchoring process, we focused on the relationship between intracellular Ca^2+^ and cell deformation during the anchoring transition. Herein, we established the detergent-treated *S. coeruleus* model (Materials and Methods and SI Appendix, Fig. S5), modified from a previously established *Blepharisma* model (41). First, *S. coeruleus* were treated with Triton X-100 in a Ca^2+^-free solution without extracellular adenosine triphosphate (ATP) to permeabilize the membrane (Fig. 4A). The cell models were 0.22 ± 0.05 mm long (mean ± SD; *n* = 51) and cone-shaped, which is usually seen in the swimming state (Fig. 4B). After demembranization above 4 × 10^−8^ M, the cell models adopted a droplet shape and their lengths decreased compared with those after demembranization in a Ca^2+^-free condition (Fig. 4B).

**Fig. 4.**
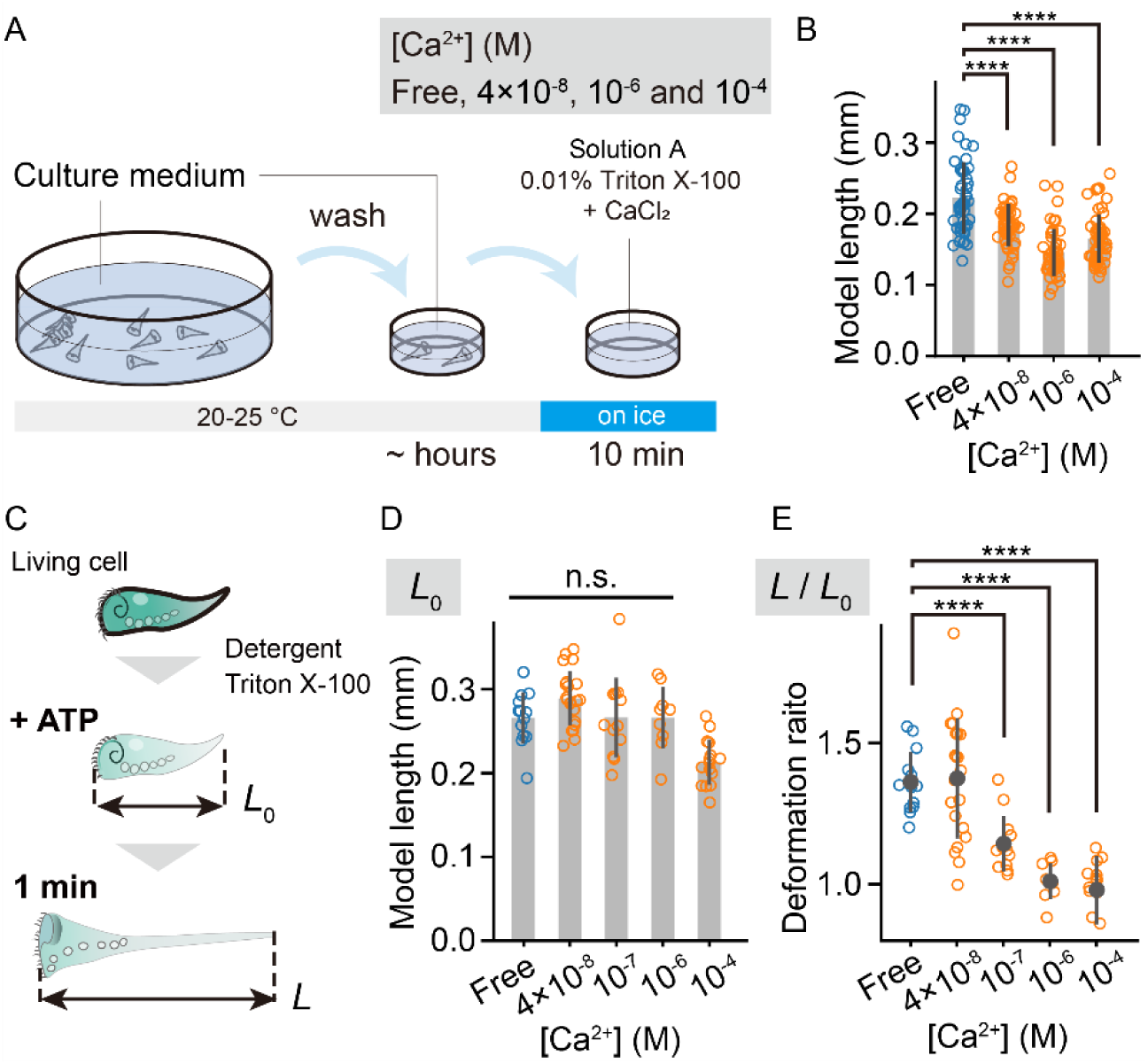
Effects of Ca^2+^ concentration on the deformations of the Triton-extracted *Stentor coeruleus* model. (A) Schematic diagram of the membrane extraction process at different Ca^2+^ concentrations. (B) High Ca^2+^ concentrations promoted contraction of the model (Welch’s *t*-test, *p* < 10^−5^, *t* = 4.62 in 4×10^−8^ M; *p* < 10^−13^, *t* = 8.95 in 10^−6^ M; *p* < 10^−8^, *t* = 6.68 in 10^−4^ M, one side, *n* = 47–51). (C) Schematics of the detergent-extracted model used to evaluate the effect of different Ca^2+^ concentrations on model elongation. (D) Initial model lengths *L*_0_ showed no significant difference at Ca^2+^ concentrations below 10^−6^ M (one-way ANOVA, *F* (3, 51) = 1.58, *p* = 0.21). The model length significantly decreased at high Ca^2+^ concentrations (10^−4^ M). (E) Deformation ratio *L* / *L*_0_ 1 min after ATP addition at different Ca^2+^ concentrations (final concentration ∼ 4 mM). For Ca^2+^ concentrations above 10^−7^ M, the elongation of the model was gradually suppressed (Welch’s *t*-test, *p* < 10^−4^, *t* = 5.11 in 10^−7^ M, *p* < 10^−8^, *t* = 9.23 in 10^−6^ M, and *p* < 10^−8^, *t* = 8.54 in 10^−4^ M, one side, *n* = 9–21). All error bars represent SDs. See also SI Appendix, Fig. S5.

The MBs and body cilia of the detergent-treated cell models exhibited beating after adding ATP at a final concentration of 4 mM in a Ca^2+^-free condition (Fig. 4C). Although some *S. coeruleus* models showed posterior elongation forming a trumpet shape, this elongation was less frequent in models with an original length of <0.25 mm (SI Appendix, Fig. S5). Thus, we excluded the droplet-shaped cell models from further analysis of Ca^2+^-dependent model elongation. Although the initial model lengths *L*_0_ did not exhibit a significant difference at Ca^2+^ concentrations of <10^−6^ M (Fig. 4D), the elongation was suppressed at Ca^2+^ concentrations of >10^−7^ M (Fig. 4E). In low Ca^2+^ concentrations, the deformation ratios 1 min after ATP addition were 1.36 ± 0.11 in the Ca^2+^-free condition and 1.38 ± 0.21 at [Ca^2+^] = 4 × 10^−8^ M. Therefore, high Ca^2+^ concentration during detergent treatment induced contraction in *S. coeruleus* models in the absence of ATP. However, model elongation required ATP and occurred at low Ca^2+^ concentration.

### Asymmetrical cell shape during transition hydrodynamically facilitates the wall-following behavior

Next, we examined how *S. coeruleus* achieved the wall-following behavior before anchoring without visual cues. *S. coeruleus* moves using cilia distributed along the entire cell surface (body cilia) and MB located around the frontal field (oral plane) at the anterior end. The water current around the MB is approximately 100 times stronger than that around the body cilia (42). We compared the orientation of the oral plane between the steady swimming and transition states (Fig. 3F and SI Appendix, Fig. S6). Oral angles were calculated from the orientation of the oral plane relative to the anterior–posterior axis. In the steady swimming state, the oral angle was slightly tilted toward the ventral side (*θ*_oral_ = 8.8° ± 5.8°, mean ± SD, *n* = 14 cells), whereas the oral angle significantly increased in the transition state (Fig. 3G, *θ*_oral_ = 30.1° ± 5.6°, mean ± SD, *n* = 11 cells). When the wall was absent on the ventral side, the cells swam in a circular trajectory, with the ventral side directed to the circular center owing to the torque induced by the asymmetric propulsion. In the steady swimming state, the asymmetric cellular shape was manifested as helical swimming with rotation around the long axis of the body.

Additionally, *S. coeruleus* without MB following sucrose shock treatment did not exhibit wall-following behavior, and cells anchored in open areas (SI Appendix, Fig. S7), indicating that MB is necessary for wall-following behavior and anchoring preference.

Based on cell deformation, we conducted a hydrodynamical simulation to investigate the mechanical relationship between the wall-following behavior and dorsoventral asymmetry of the oral plane. To construct the numerical *Stentor*, we defined point forces that were distributed only around the base of a conical object (Fig. 5A and SI Appendix, Fig. S8). The model excluded both body cilia activity and the effect of rotation around the long axis. Assuming force-and torque-free constraints arising from viscous fluid dynamics, the swimmer’s translational and angular velocities were determined at each time step using a boundary element method. Based on these values, the swimmer’s centroid position and orientation were calculated as time series. The elongated portion of the swimmer was horizontally displaced parallel to the base of the conical object, which constrains the swimmer’s volume (Fig. 5B). While the symmetrical numerical *Stentor* approached the wall and then deflected away owing to the upward torque generated by hydrodynamic interactions (Fig. 5C), the numerical *Stentor* with asymmetrical propulsion turned toward the wall owing to self-generated downward torque and moved along the wall (Fig. 5D and Movie S4). When the downward torque caused by the asymmetrical distribution of propulsive forces exceeds the upward torque generated hydrodynamically, the net torque becomes zero, resulting from a direct repulsive force from the substrate, enabling the swimmer to move along the wall. With small asymmetricity (*θ*_oral_ = 5°–15°), the swimmer temporarily swam away from the wall owing to the near-wall hydrodynamic interactions and returned to the wall using self-generated downward torque, resulting in two motions repeatedly and causing wave-like trajectories (Fig. 5E). The oral angles in the wave-like trajectories of the simulation were rare in the experimental transition state (Fig. 3G). Therefore, the dorsoventral asymmetry of propulsion without rotation along the long axis of the cell provides a physical explanation for the wall-following behavior of *S. coeruleus*.

**Fig. 5.**
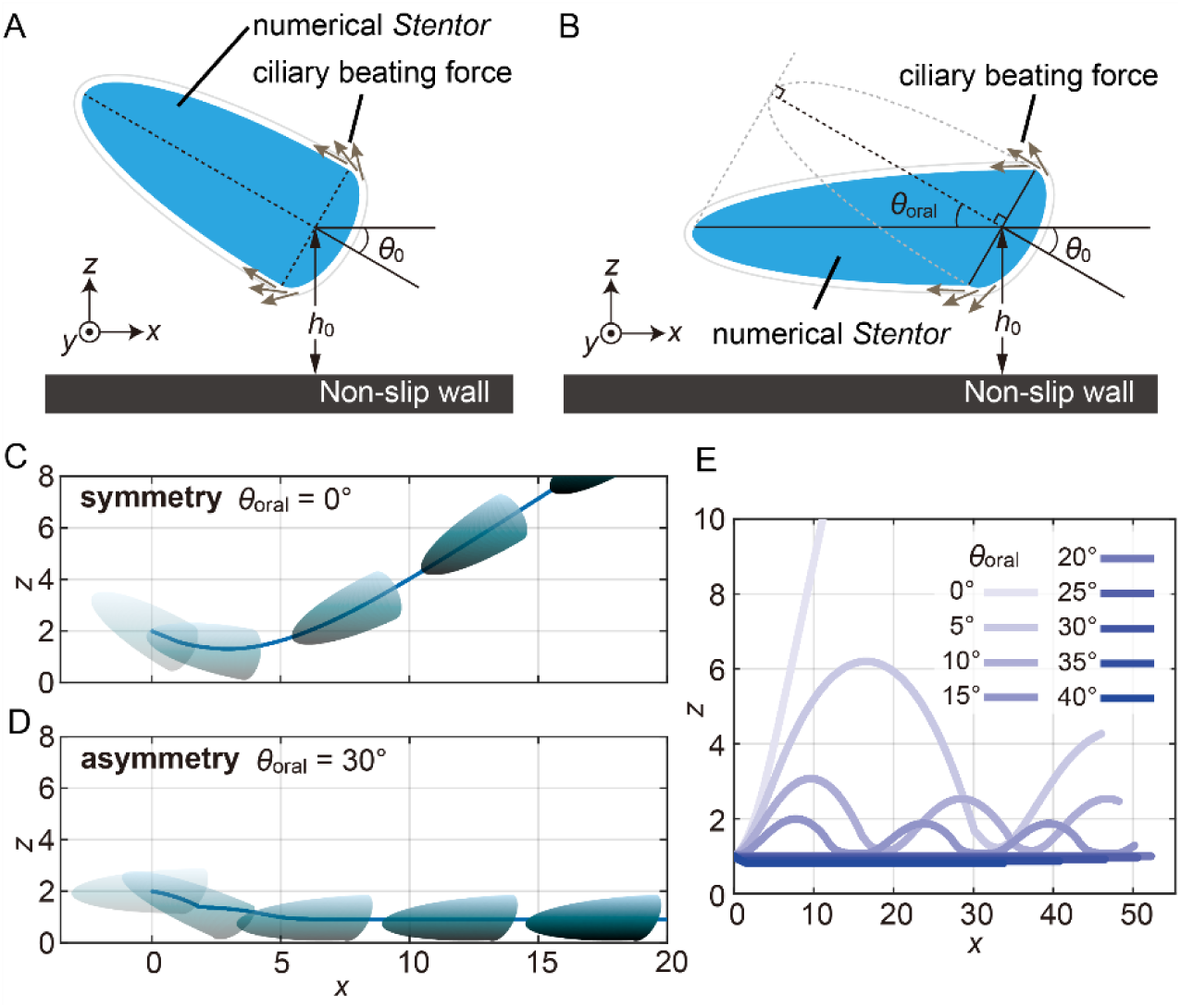
Asymmetric propulsion of a conical swimmer physically explaining the wall-following swimming behavior. **(A)** Schematic of the numerical simulation. A conical object, representing a simplified *Stentor* cell shape, deformed from a sphere of radius 1. The object at (*x*, *y*, *z*) = (0, 0, 2) in the initial state faces the boundary wall at *z* = 0. Point forces, which correspond to the thrust forces from the beating MB, are distributed peripherally around the base of the conical object tangency, directing the base to the vertex. **(B)** Schematic of the deformed swimmer. The elongated half of the swimmer horizontally shifts along the contracted hemisphere surface of the conical object (A). The initial condition and distribution in force are the same as (A). **(C, D)** Time evolution and trajectories of symmetrical and asymmetrical swimmers. Each center of the swimmer is initially located at (0, 0, 2), and the locations of the conical swimmer are shown at 1, 200, 800, 1400, and 2000 numerical time steps. Although the symmetrical conical swimmer reflects the boundary owing to hydrodynamic interactions (C), the asymmetrical conical swimmer moves along the wall (D). **(E)** Trajectories of the swimmer depending on the asymmetricity of the oral angle. Each center of the swimmer is initially located at (0, 0, 1), and the posterior axis horizontally aligns with the wall. See also SI Appendix, Fig. S8 and Movie S4.

### Different interactions between wall-following *S. coeruleus* and the boundary wall depending on geometrical features

To address the selection of anchoring sites formed by different geometric boundaries, we examined the swimming speed of wall-following *S. coeruleus* in geometrically structured chambers as a function of the distance from the corner or channel tips. Figure 6 shows that swimming speed tended to decrease near the corner and channel tips. In right triangle chambers, the speed near corners with smaller angles decreased relative to that near larger angles (Fig. 6A). As shown in Fig. 6B, although a similar tendency was observed in out-of-corner structures, swimming speed decreased around acute corner regions, particularly at 30° and 45° corners, where the cells frequently anchored. Distinct contact interactions were observed, accompanied by transient suppression of movement around the 30° corner (SI Appendix, Fig. S9 and Movie S7).

**Fig. 6.**
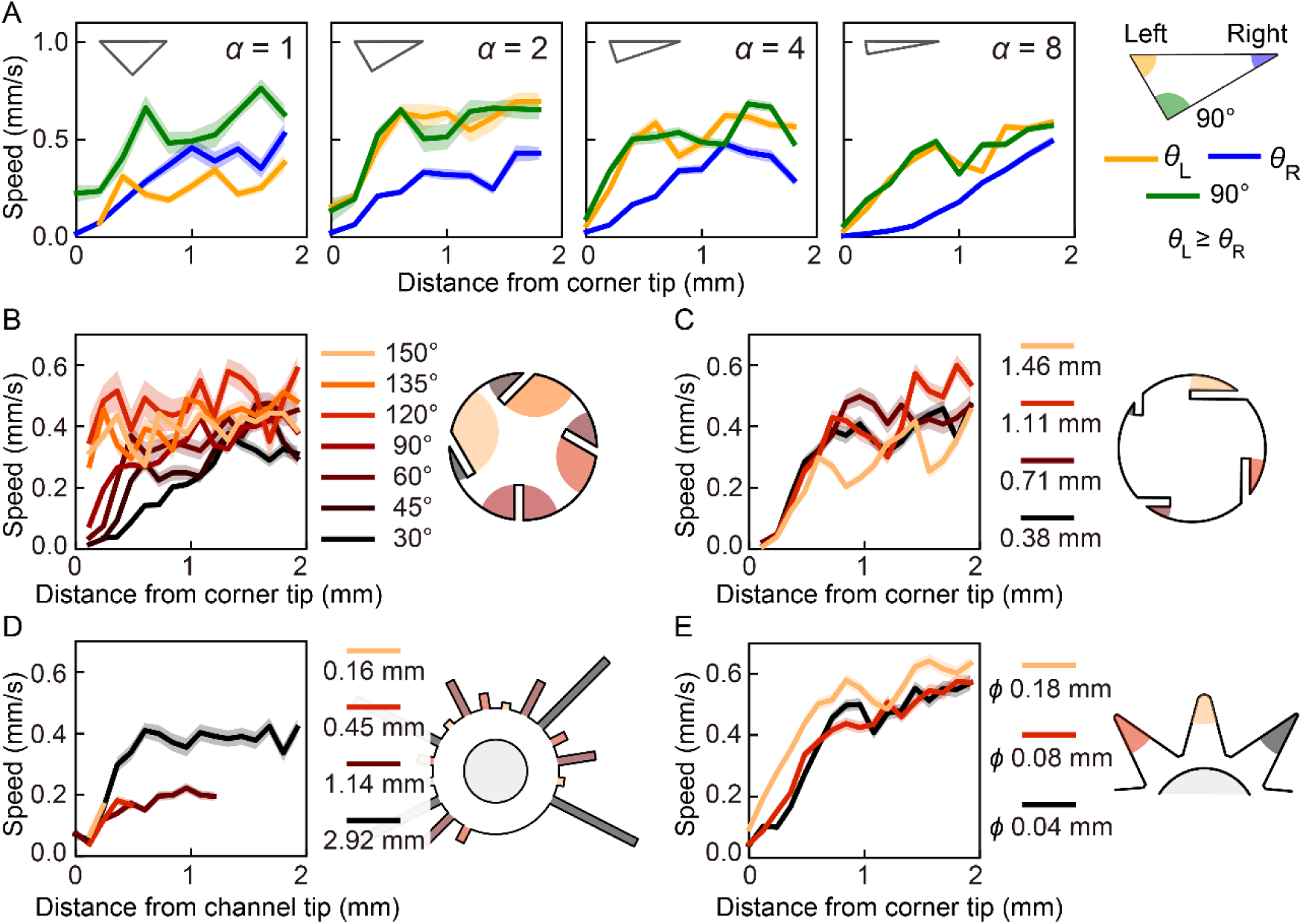
Swimming speed of wall-following *S. coeruleus* as influenced by surrounding geometrical features. The distribution of averaged swimming speed was calculated in geometrically structured chambers as a function of distance from the corner or channel tip. Solid lines and shaded bands represent mean values and standard errors, respectively. Line colors correspond to the colored regions in the schematic illustration of the chamber boundary shown on the right. Observation chambers (A) to (E) correspond to Fig. 2D–H. See also SI Appendix, Figs. S9, S10 and Movies S5–S9.

Similar tendencies in swimming speed were observed near tips with respect to corner and channel depth (Fig. 6C, D). This indicates that the cells could anchor near short protrusion structures owing to deceleration in swimming speed. Although swimming speed decreased at the tip of the shallowest channel, the cell could not remain within the channel because its depth was shorter than the cell length. Additionally, swimming speed was not suppressed beyond the channel tip (distance > 0.5 mm) at a depth of 2.92 mm (Fig. 6D), suggesting no facilitation of anchoring.

Deceleration was also observed near corners with low curvature radii of 0.04 mm and 0.08 mm compared with swimming speed near round corners (curvature radius = 0.18 mm) (Fig. 6E and SI Appendix, Fig. S10, Movies S8 and S9). From these results, although some exceptions were observed, the swimming speed of wall-following *S. coeruleus* decreased near corner tips with geometrical features where *S. coeruleus* preferentially anchors.

## Discussion

We quantified the detailed geometrical characteristics of narrow areas influencing the anchoring. The anchoring processes from swimming to adhering involved wall-following exploration with asymmetrical morphological change associated with intracellular Ca^2+^ concentrations. A mechanical model and observation of the cell deformation indicated that asymmetric propulsion facilitates the wall-following behavior not through continuous visual recognition, enabling final anchoring to narrow sites (Fig. 7).

**Fig. 7.**
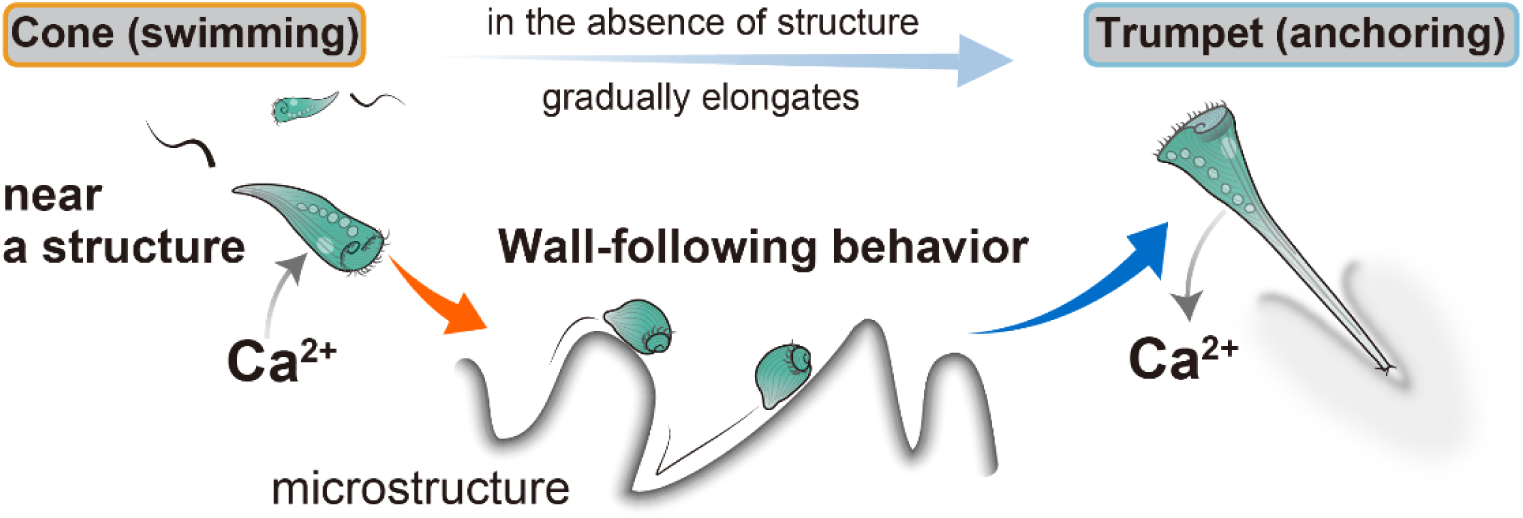
Schematic illustration of the anchoring behavior in the presence of structures around the cell. *S. coeruleus* usually swims with a helical trajectory during the steady swimming state. In the presence of nearby structures, frequent contact with the wall causes Ca^2+^ influx, triggering gradual cell contraction, and the cell exhibits an asymmetric propulsive force, physically inducing a wall-following behavior. Finally, the movement is physically regulated by the surrounding geometrical structures, and the deceleration facilitates anchoring. During the elongation process, Ca^2+^ efflux occurs. In contrast, in the absence of a structure, the cell gradually elongates from a conical swimming shape to a trumpet shape without contraction.

The wall-following behavior was observed only in geometrically complex environments, indicating that an increase in contact with a wall due to the geometrical complexity induces the anchoring transition. Ciliates referred to as “swimming neuron” (43, 44) typically exhibit transient backward swimming upon collision against a wall, a response triggered by the bioelectric activity of the membrane potential through Ca^2+^ influx (33, 45–47). Consistent with this, our contracted permeabilized *Stentor* model with high Ca^2+^ concentration exhibited contraction, supporting the idea that increased contact triggers contraction. Furthermore, backward swimming and anchoring were positively correlated in space (Fig. 2D, E, H; SI Appendix, Fig. S3), and mechanical contact events were observed immediately before wall-following behavior (SI Appendix, Fig. S11, Movies S10 and S11). Given the frequent contacts in confined observation spaces, these observations suggest that repeated mechanical stimuli can drive *S. coeruleus* to transition into wall-following behavior.

If the contractile fiber myoneme is asymmetrically distributed, as suggested (48, 49), contraction may tilt the oral plane, thereby contributing to the asymmetric morphological change causing wall-following behavior. The wall-following *S. coeruleus* that facilitated entry into corners interacted differently with contacting wall boundaries depending on boundary shape. Swimming speed markedly decreased near corner tips where *S. coeruleus* preferentially anchored (SI Appendix, Figs. S9, S10, Movies S7 and S9). In contrast, *S. coeruleus* escaped regions along the wall where anchoring did not occur (Movies S5, S6 and S8). The geometries where cells frequently anchored corresponded to cellular dimensions such as length and width. This suggests that deceleration of swimming speed owing to direct contact with the wall in specific regions facilitates adherence at these locations. Overall, anchoring preference emerged from physical interactions between the embodied organism and the geometrical wall during wall-following behavior caused by intrinsic cellular deformation.

Animals that rely on visual cues utilize surrounding geometrical structures for navigation (6, 9). *Stentor* is a well-studied model system for behavioral complexity at the single-cell level, including habituation (50, 51) and hierarchical responses (52, 53) to repeated mechanical stimuli. Electrophysiological studies have provided mechanistic insights into stimulus-dependent changes in responsiveness (54, 55). We found that *S. coeruleus*, despite lacking visual cues, also exhibits anchoring-site preference based on surrounding geometry. These findings are expected to advance our understanding of how unicellular organisms differentially sense a range of mechanical stimuli.

Although wall-following behavior is conserved across several organisms from unicellular species to animals (56), the wall-following behavior observed in *S. coeruleus* exhibits two distinct biophysical mechanisms. First, while hydrodynamic interactions of microswimmers with the wall have been widely investigated (57–59), our findings reveal a novel and simple hydrodynamic mechanism: asymmetric propulsive forces drive the thigmotactic behavior in a puller-type microswimmer that possesses a propulsion apparatus located at the front of the body (60). This mechanism has the potential to describe a broad range of thigmotactic behaviors in puller-type swimmers. Second, although myoneme-induced rapid contractions in ciliates typically occur in milliseconds (61), the contraction duration observed in our study (Fig. 3A) was approximately 1000 times longer than that of a rapid contraction. The contraction during the anchoring process in *Stentor* has not been reported.

Anchoring in narrower sites of geometrically structured chambers may help avoid predators, light exposure, and desiccation. Large predators rarely enter into narrow areas, which are covered by structures reducing light. Furthermore, under drying conditions, water is likely to accumulate in narrow areas owing to surface tension. Our results revealed rapid contraction in *Stentor*, likely to avoid predators, allowing the entire cell body to fit within the structural narrowness. Moreover, *Stentor* preys on swimming algae, *Chlamydomonas*, which aggregates at corners due to physical mechanisms (31). *Stentor* might be able to obtain the food despite being in stagnant narrow areas. Therefore, the adaptive behavioral transition of *S. coeruleus* observed in geometrically complex environments can be mechanically explained by the simple asymmetrical deformation of the cell.

In conclusion, we identified the geometrical features of the preferential anchoring sites of *S. coeruleus* using geometrically structured chambers. The anchoring process involves intrinsic cellular deformation causing the wall-following and contact interaction. A simple mechanism of the asymmetric cell contraction physically explains the wall-following behavior. *S. coeruleus* anchors due to a decrease in motility caused by the surrounding geometrical features. Our findings provide evidence that even in the absence of a visual input, the unicellular organism ubiquitously utilizes microgeometrical features in its habitats. Diverse geometrical shapes in microhabitats have the potential to create ecological niches for protists (62, 63), including *Stentor*, which play crucial roles in food chains (64, 65). Behavioral self-organization of aquatic organisms can influence ecosystem dynamics (66, 67). Together, these findings offer insights into primitive navigation systems in unicellular organisms and highlight the relationship between environmental geometry and ecological networks.

## Materials and Methods

### Cell culture

*S. coeruleus* were collected from the Shiribetsu River (42.81°N, 140.69°W) in Japan. The cultures were maintained using modified Peters’ solution (68) containing an oat grain. Details are provided in the SI Appendix.

### Observation of the anchoring behavior in geometrically structured chambers

Geometrical observation chambers were fabricated via a molding process. These molds were designed using the three-dimensional (3D) CAD software Fusion360 (Autodesk, California, United States) (SI Appendix, Fig. S1). A milling machine, CAMM-3 Model PNC-3200 (Roland DG Corporation, Hamamatsu, Japan), equipped with a microend mill was used to fabricate molds by cutting the modeling wax ZW-100 (Roland DG Corporation, Hamamatsu, Japan). Prior to making the geometrically structured chambers, 2.0% (w/v) agarose solution was prepared from the modified Peters’ solution and PrimeGel (Agarose LE 1–20K, Takara Bio, Shiga, Japan). After the mixture was melted, it was poured into the mold and sealed with a glass slide to flatten the upper surface (SI Appendix, Fig. S1). After solidification, the gel chamber was detached from the mold.

Prior to observation, cells were transferred into fresh culture medium twice to prevent chemical effects, allowing the cells to acclimate for several hours. The geometrically structured chambers were rinsed with fresh medium on a glass slide, and the chamber was filled with the medium and sealed with a coverslip pretreated with oxygen plasma for 10–20 s to increase surface hydrophilicity (PC-400T, STREX, Osaka, Japan). An individual swimming *S. coeruleus* was gently released into the gap between the chamber and the above coverslip. Subsequently, the observation chamber was entirely sealed by sliding the coverslip into place. The behaviors were recorded using a CMOS camera, ORCA-spark C11440-36U (Hamamatsu Photonics, Hamamatsu, Japan) mounted on a stereo microscope, SZX16 (Evident, Tokyo, Japan). The images were acquired at 4 frames per second (fps) with a 50 ms exposure time. Imaging was conducted under red-filtered brightfield illumination and minimal light intensity to prevent phototactic responses in *S. coeruleus*.

### Quantification of geometrical preferences of anchoring sites

Anchoring sites and durations were manually obtained, and anchoring events shorter than 10 s were excluded. Anchoring site preferences in the Results section were quantified using two metrics: 1) average anchoring rate within each defined region, which included a characteristic geometrical feature formed by an outer wall of the geometrically structured chamber and 2) probability density function (PDF) of anchoring, defined as the average anchoring rate divided by the area of each region. Detailed definitions of each region are provided in the SI Appendix (Fig. S1).

To evaluate the effect of corner angle on anchoring in right triangle chambers, an asymmetry index was calculated from the anchoring rate in left *θ*_L_ or right *θ*_R_ angles per area when facing the 90-degree corner (*θ*_L_ ≥ *θ*_R_), as defined below.

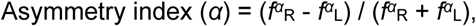

where *α* is a shape parameter in triangle chambers defined as *θ*_L_/*θ*_R_. The PDF of anchoring in the *α*-shaped chamber, *f^α^*_i_ is defined as

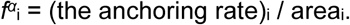

The geometrical values reported in the Results section were measured from the actual agarose chambers, rather than from the design.

### Image analysis and quantification of behaviors in the geometrically structured chambers

Cell center and body length were extracted from image contours using ImageJ/Fiji (69) and Python 3.11.7 (https://www.python.org/). The recorded images were subtracted using a background image acquired from sequential frames. Gaussian blur was applied to the background-subtracted sequential images to reduce noise. Then, the images were binarized by applying the brightness thresholds. Cell length was calculated from the binary images as the sum of two distances measured from the center of the cell to its anterior and posterior ends (32). Since Gaussian blur caused overestimation of the body length by 20–100 μm, we corrected the measured lengths by comparing the cell outline and raw data. Swimming velocity was calculated from the displacement of the center of the cell. Both velocity and cell length data were smoothed by the median values of ± 10 s.

### Comparison of anchoring transition with and without geometrical structures

To compare anchoring behaviors in the presence or absence of geometrical structures, we recorded individual behavior in a simple circular chamber using a CMOS camera, ORCA-Flash4.0 v2 (Hamamatsu Photonics, Hamamatsu, Japan), under red-filtered brightfield illumination with an exposure time of 50 ms at 4 fps with minimal light intensity. A simple circular chamber was fabricated using agarose gel into a quasi-two-dimensional disk of 0.3 mm thickness and 5 mm diameter.

We defined the start of the anchoring transition as the point when the cell deformed from its steady cone-shaped swimming form, and the end as the point when the cell became sufficiently elongated, indicating the final state. The cell length over time was normalized to the individual mean cell length in the steady-swimming state. We classified the behavioral transition as “contraction” when the mean length 30 s after deformation was shorter than that in the steady swimming state, and as “elongation” when it was longer.

To compare swimming trajectories during steady swimming and the transition before anchoring, we manually selected a helical steady-swimming phase and the transition section (each up to 1 min). The transition section was defined to start 15 s after cell deformation, excluding any period when the cell was anchored. The distance between the cell and the chamber wall was measured as the minimum distance between the cell center and the outer wall. Additionally, we quantified the dwell time of the cell within 0.15 mm of the outer wall to evaluate contact duration with the wall.

In these analyses, we calculated the mean values for each cell over its recording period and then calculated the mean and standard deviation (SD) for each population.

### Measurement of the orientation angle of the oral plane

To determine the oral structure, we obtained high-magnification cell images using the same experimental setup in the geometrically structured chamber (SI Appendix, Fig. S6A). A 1-ms exposure time in these observations allowed visualization of the ciliary band (MB). We selected one representative image to quantify the cellular shape in the steady swimming state and the wall-following state, with two body axes, anterior–posterior (AP) and dorsoventral (DV) (70), aligned to an image plane because the swimming *S. coeruleus* rotated along the AP axis. An oral plane is formed by a closed area of the MB located at the periphery of the oral apparatus at the anterior end. Owing to the three-dimensional structure of the oral plane resulting from misalignment of circular MB on the ventral side, we manually selected two points on the ventral side and one point on the dorsal side of the MB, where light intensity exceeded background levels (SI Appendix, Fig. S6B). From the two segments connecting the ventral and dorsal points, we calculated an averaged segment and determined the tilt angle of the oral plane relative to the DV axis, defining the ventral side as the positive direction (SI Appendix, Fig. S6C, D). The AP axis was defined as the line connecting the posterior end to the midpoint of the averaged segment. Detailed methods are provided in the SI Appendix, Fig. S6.

### Development of a detergent-treated model of *S. coeruleus* and observation methods

We developed a detergent-treated *S. coeruleus* model modifying a cell model in *Blepharisma*, which is closely related to *Stentor*, as described in Matsuoka et al. 1991 (41). The detailed protocol, solution composition and observation conditions are provided in the SI Appendix.

### Measurement of model length and evaluation of elongation

We manually segmented three points from the anterior to the posterior end and determined the model length as the total length of the two segmented lines (SI Appendix, Fig. S5). The elongation ratio was calculated by the initial length *L*_0_, measured within 30 s after ATP addition, and the length *L,* measured 1 min after ATP addition. If accurate measurement of the length at the 1-min time point was not possible due to three-dimensional motion, the nearest available time frame was used.

### Construction of a hydrodynamical model of a numerical *Stentor*

To investigate the movement of the numerical swimmer near the wall, we developed a conical swimmer model mimicking *Stentor*’s shape by deforming a spherical object with radius 1. Point forces mimicking ciliary beating were distributed around the base of the conical swimmer. The deformation method, propulsive force distribution and initial conditions for numerical simulations are described in the SI Appendix.

### Computational simulation of movement in the numerical swimmer near a plane boundary

We computed the motion of the numerical swimmer under force-and torque-free constraints arising from the nature of viscous fluids using a boundary element method, following the scheme described by Ohmura et al. (26). Detailed numerical methods are provided in the SI Appendix.

### The interaction of wall-following *S. coeruleus* with different boundary geometries

From our experimental recordings, we manually selected periods during which *S. coeruleus* was in the wall-following state, defined by characteristic nonhelical swimming motion with a droplet-like cell shape. We then analyzed swimming speed as a function of distance from corner or channel tips for each geometrical feature. The final 15 seconds of each period were excluded from the analysis because motionless *S. coeruleus* did not interact with the boundary wall immediately before anchoring. Averaged swimming speed and standard error were calculated for each 0.06–0.1 mm interval relative to distance from the tips.

### Quantification and statistical analysis

Statistical analyses were conducted using scipy.stats version 1.11.4 in Python 3.11.7. Depending on the comparison, paired *t*-tests, Welch’s *t*-tests, and one-way ANOVA were used. Test statistics and sample sizes are provided in the Figure legends and main text. Statistical significance was indicated as follows: n.s. (not significant, *p* > 0.05); \*\**p* < 0.01; \*\*\**p* < 0.001; \*\*\*\**p* < 0.0001.

## Acknowledgments

We would like to thank Takuji Ishikawa and Kenta Ishimoto for their useful discussions and comments on the theoretical aspects. We also thank Osamu Kishida, Norihiko Kamaya and Alid Al-Asmar for collecting samples and discussing the ecological aspects. Taku Morimoto and the Division of Technical Staffs in the Research Institute for Electronic Science at Hokkaido University supported the fabrication technique.

## Author Contributions

Conceptualization: SE, KS, TN, YN; Methodology: SE, TN, YN; Software: TO; Formal Analysis: SE; Investigation: SE, TO; Resources: SE, TO, YN; Data Curation: SE, TO; Writing – Original Draft: SE; Writing – Review & Editing: SE, TO, KS, TN, YN; Visualization: SE, TO; Supervision: TN, YN; Funding Acquisition: SE, TO, KS, TN, YN

## Data and materials availability

All data needed to evaluate the conclusions in the paper are present in the paper and/or the Supporting Information.

