## Supplementary Information for "Geometrical preference of anchoring sites in the unicellular organism *Stentor coeruleus*"

### **Supplementary Information for**
### Geometrical preference of anchoring sites in the unicellular organism *Stentor coeruleus*

Syun Echigoya[1,2]*, Takuya Ohmura[1,3], Katsuhiko Sato[4], Toshiyuki Nakagaki[1,3], Yukinori Nishigami[1,3]*

**This PDF file includes:**

- Supporting Information text
- Figures S1 to S11
- Legends for Movies S1 to S11
- SI References

#### Cell culture

*S. coeruleus* were collected from the Shiribetsu River (42.81°N, 140.69°W) in Japan. The cultures were maintained in a 6-cm plastic dish filled with modified Peters' solution (1) (0.55 mM $CaCl_2$, 0.15 mM $MgSO_4$, 0.15 mM $K_2CO_3$, and 0.75 mM $Na_2CO_3$, adjusted pH 7.4 ± 0.1 with 1 N HCl) containing an oat grain (MRP-706, Marukan, Osaka) to promote bacterial growth as a nutrient at 25 ℃ in dark condition. The cells were subcultured into fresh medium once they reached a high cell density, typically every 2–3 weeks.

#### Definition of analyzed regions of anchoring sites in geometrically structured chambers

To evaluate the site preference of anchoring in geometrically structured chambers, we defined the regions for analysis (Fig. S1A–E). The regions analyzed in right triangle chambers were defined as corners within 1 mm from the corner tips. Overlapping regions with two corners were divided and classified into the nearest corner. Regions analyzed in the Angle-chamber and the Depth-chamber were the inner regions between the protrusion and the outer wall, within each protrusion length. The regions analyzed in the Channel-chamber were the inner regions of each channel. The regions in the Star-chamber were within $L' = L$ + offset (mm) from the intersection of the outer walls in each corner, where $L = 2$ and offsets represent the manually measured distances from the wall intersection to each corner tip. The two-dimensional surface areas of each region were measured from CAD designs of the geometrically structured chambers rescaled to the original size.

#### Development of a detergent-treated model of *S. coeruleus* and observation methods

We developed a detergent-treated *S. coeruleus* model modifying a cell model in *Blepharisma*, which is closely related to *Stentor*, as described in Matsuoka et al. (2). Prior to the experiment, *S. coeruleus* were transferred into a fresh modified Peters' solution to remove other chemicals and allow acclimatization for several hours under dim light. To investigate the influence of $Ca^{2+}$ concentration on the cell deformation without ATP, 10–20 cells were immersed in a detergent medium, solution A (20 mM 1,4-Piperazinediethanesulfonic acid (PIPES) (pH = 6.9, adjusted by KOH), 5 mM ethylene glycol tetraacetic acid (EGTA) (pH = 7.0, adjusted by KOH), 50 mM KCl, 4 mM $MgSO_4$), and 0.01% (v/v) Triton X-100) with $CaCl_2$ adjusting $Ca^{2+}$ concentration from $4 \times 10^{-8}$ M to $10^{-4}$ M or without $Ca^{2+}$ in a plastic dish on ice (Fig 4A). The dish was gently agitated intermittently to remove the surface layer of cells from the model until the MB ceased beating completely. This process was recorded using a CMOS camera, ORCA-Flash4.0 v2, mounted on stereomicroscope, SZX16, under brightfield illumination with 5 ms exposure time, 2x2 binning, and 1 fps.

To measure the effect of $Ca^{2+}$ concentration on the ATP-induced elongation, detergent-treated cell models were prepared in $Ca^{2+}$-free solution and then transferred to a plastic dish containing solution A on ice to remove residual detergent. Subsequently, they were treated at 20 ℃–25 ℃ for 1 min in solution A with $CaCl_2$ from $4 \times 10^{-8}$ M to $10^{-4}$ M or under $Ca^{2+}$-free conditions. After ATP addition (final concentration of 4 mM), the model deformation was observed on a glass slip enclosed by a 0.5 mm-thick silicon sheet with a 5 mm inner diameter well (Fig. S5A). The solution volume was adjusted so that the top meniscus remained as horizontal as possible. Imaging was performed using a CMOS camera, ORCA-Flash4.0 v2, mounted to a stereomicroscope, SZX16, under brightfield illumination with 20 ms exposure time and 1x1 binning. The cell models with irregular shapes were excluded. Moreover, since droplet-shaped short models did not exhibit any elongation (Fig. S5C–F), they were excluded to focus solely on models capable of elongation. During cell transfers, plastic pipette tips were coated with Lipidure® CM5206 to prevent cell adhesion. The free $Ca^{2+}$ concentration was determined using the Ca-Mg-ATP-EGTA Calculator v1.0 (https://somapp.ucdmc.ucdavis.edu/pharmacology/bers/maxchelator/CaMgATPEGTA-NIST.htm) based on the constants from NIST database #46 v8, under pH 7.0 and ionic strength of 0.1 N.

#### Detailed explanation of measurement of the orientation angle of the oral plane

To quantify the extent to which the oral plane formed by MB tilts toward the ventral side, high-magnification observations with a 1-ms exposure time were performed to visualize the ciliary band (MB). *S. coeruleus* exhibits distinct cellular structures, with a flat ventral side and a gently convex dorsal side (3). Figure S6A shows sequential side-view images of *S. coeruleus*. For analysis, we selected an image in which the anterior–posterior and the ventral–dorsal axes were aligned with the image plane (indicated by the orange solid rectangle in Fig. S6B). The tips of MB around the oral apparatus are discontinuous on the ventral side (3). Due to the three-dimensional warped structure, we selected three points on the MB based on regions of higher brightness in the captured image. Specifically, two points, S and R, were selected on the ventral side, and one point, Q, on the dorsal side was selected as the midpoint of the MB perimeter. Point M was defined as the midpoint of line segment SR, and an averaged segment was defined by QM. Point A was defined as the midpoint of segment QM, and the anterior-posterior (A–P) axis was defined by segment AP, where point P corresponds to the posterior endpoint. The ventral–dorsal (V–D) axis was defined by line VD, orthogonal to line AP. Finally, the oral angle was defined as the angle between VD and QM.

#### Construction of a hydrodynamical model of a numerical *Stentor*

We developed a conical swimmer model mimicking *Stentor*'s shape while swimming, by deforming a spherical object with radius 1, achieved by linearly compressing a hemisphere and linearly expanding another hemisphere (Fig. S8). We considered propulsive forces only around the anterior end, excluding the propulsion of body cilia, because a swimming current around the anterior MB is tens of times stronger than that around the entire cell body. Point forces mimicking ciliary beating were distributed around the base of the conical swimmer. The beating direction was assumed to be tangential to the body surface oriented from the base to the vertex of the conical object. The orientation angle of the oral plane ($\theta_{\mathrm{oral}}$) was modulated by linearly sliding the hemisphere, whose elongated side is horizontally against the base, conserving the swimmer's volume. $\theta_{\mathrm{oral}}$ represents the angle between the axes of the two deformed hemispheres. We numerically solved the translational velocity $\boldsymbol{U}$ and angular velocity $\boldsymbol{\Omega}$ of the swimmer from the center of mass for $\theta_{\mathrm{oral}} = 0$ or 30° in a 3D viscous fluid on a nonslip boundary located at $z = 0$ with the initial location of the center of swimmer's mass $\boldsymbol{r}_g = (0, 0, 2)$ and the initial incidence angle between the direction of the anterior side and the nonslip boundary $\theta_0 = 30°$. To investigate the movement of the numerical swimmer near the wall, we set the initial conditions as $\boldsymbol{r}_g = (0, 0, 1)$ and $\theta_0 = \theta_{\mathrm{oral}}$, which means that the angle between the posterior axis and the boundary is zero. Then, swimming trajectories were obtained from the calculated $\boldsymbol{U}$ and $\boldsymbol{\Omega}$.

#### Computational simulation of movement in the numerical swimmer near a plane boundary

For swimming dynamics for *Stentor*, viscous effects are more dominant than inertia (Reynolds number, $Re \sim 0.3$), which allows us to describe the swimming behavior using low-Reynolds-number hydrodynamics. To calculate hydrodynamics around the numerical swimmer at low Reynolds number, a boundary element method (BEM) in viscous flow was employed (4). For calculating the motion of the numerical swimmer defined in the above section, we followed the scheme used in Ohmura et al. (5).

In a viscous fluid, the swimming speed $\boldsymbol{U}$ and angular velocity $\boldsymbol{\Omega}$ are determined by the balance of the total force and torque from canceling propulsive forces by viscous drag. Moreover, nonslip boundary conditions were assumed on the surface of the swimmer and the boundary of the wall. Therefore, we calculated the swimming speed and angular velocity under two boundary conditions: 1) the force and torque balances between the propulsive force by swimmer $\boldsymbol{F}^{\mathrm{cilia}}(\boldsymbol{r}'')$ and the stress of viscous drag $\boldsymbol{q}^{\mathrm{body}}(\boldsymbol{r}')$

$$\int_{\mathrm{cilia}} \boldsymbol{F}^{\mathrm{cilia}}(\boldsymbol{r}'')\mathrm{d}S(\boldsymbol{r}'') = \int_{\mathrm{swimmer}} \boldsymbol{q}^{\mathrm{body}}(\boldsymbol{r}')\mathrm{d}S(\boldsymbol{r}'),$$

$$\int_{\text{cilia}} \boldsymbol{F}^{\text{cilia}}(\boldsymbol{r}'') \times (\boldsymbol{r}'' - \boldsymbol{r}'_{\text{g}}) \text{d}S(\boldsymbol{r}'') = \int_{\text{swimmer}} \boldsymbol{q}^{\text{body}}(\boldsymbol{r}') \times (\boldsymbol{r}' - \boldsymbol{r}'_{\text{g}}) \text{d}S(\boldsymbol{r}')$$

equation (1),

and 2) the velocity field on the swimmer surface $\boldsymbol{u}_{\text{s}}(\boldsymbol{r}')$ caused by the propulsion satisfies the nonslip boundary conditions on the swimmer surface moving at $\boldsymbol{U}$ and $\boldsymbol{\Omega}$ around the center of mass $\boldsymbol{r}_{\text{g}}$ in a static fluid

$$\boldsymbol{u}_{\text{s}}(\boldsymbol{r}') = \boldsymbol{U} + \boldsymbol{\Omega} \times (\boldsymbol{r}' - \boldsymbol{r}'_{\text{g}})$$

equation (2),

where $\boldsymbol{r}'$ is a location on the swimmer surface. The flow field $\boldsymbol{u}(\boldsymbol{r})$ in an infinite viscous fluid is determined from the surrounding force distribution $\boldsymbol{q}^{\text{body}}(\boldsymbol{r}')$ and $\boldsymbol{F}^{\text{cilia}}(\boldsymbol{r}'')$ by following the boundary element integration

$$u_{\text{i}}(\boldsymbol{r}) = u_{\text{i}}^{\infty} - \frac{1}{8\pi\eta}\int_{\text{swimmer}} J_{\text{ij}}\,(\boldsymbol{r},\boldsymbol{r}')\, q_{\text{j}}^{\text{body}}\,(\boldsymbol{r}')\,\text{dS}\,(\boldsymbol{r}') + \frac{1}{8\pi\eta}\int_{\text{cilia}} J_{\text{ij}}\,(\boldsymbol{r},\boldsymbol{r}'')\, F_{\text{j}}^{\text{cilia}}\,(\boldsymbol{r}'')\,\text{d}S\,(\boldsymbol{r}''),$$

$$J_{\text{ij}}(\boldsymbol{x},\boldsymbol{y}) = J_{\text{ij}}^{\text{O}}(\boldsymbol{x},\boldsymbol{y}) = \frac{\delta_{\text{ij}}}{s} + \frac{s_{\text{i}}s_{\text{j}}}{s^3},$$

$$\boldsymbol{s} = s_{\text{i}} = \boldsymbol{x} - \boldsymbol{y}, \qquad s = |\boldsymbol{s}|, \qquad u(\boldsymbol{r}) = (u_{\text{x}}(\boldsymbol{r}), u_{\text{y}}(\boldsymbol{r}), u_{\text{z}}(\boldsymbol{r}))$$

equation (3),

where $\boldsymbol{u}^{\infty}$ is the background flow velocity at infinity, $\boldsymbol{x}$ is the measurement point, and $\boldsymbol{y}$ is the location of the applied force. In this simulation, $\boldsymbol{u}^{\infty} = \boldsymbol{0}$. In a viscous fluid on an infinite nonslip plane, we can solve the flow velocity by substituting $J_{\text{ij}}$ in equation (3) into the following Blakelet tensor $J_{\text{ij}}^{\text{Blake}}$ (6),

$$J_{\text{ij}}(\boldsymbol{x},\boldsymbol{y}) = J_{\text{ij}}^{\text{Blake}}(\boldsymbol{x},\boldsymbol{y}) = J_{\text{ij}}^{\text{O}}(\boldsymbol{x},\boldsymbol{y}) - J_{\text{ij}}^{\text{O}}(\boldsymbol{x},\overline{\boldsymbol{y}}) - 2h^2 J_{\text{ij}}^{\text{SD}}(\boldsymbol{x},\overline{\boldsymbol{y}}) + 2hJ_{\text{ij}}^{\text{FD}}(\boldsymbol{x},\overline{\boldsymbol{y}}),$$

$$J_{\text{ij}}^{\text{SD}}(\boldsymbol{x},\overline{\boldsymbol{y}}) = (1 - 2\delta_{\text{jz}})\left(\frac{\delta_{\text{ij}}}{R^3} - \frac{3R_{\text{i}}R_{\text{j}}}{R^5}\right),$$

$$J_{\text{ij}}^{\text{FD}}(\boldsymbol{x},\overline{\boldsymbol{y}}) = (1 - 2\delta_{\text{jz}})\left(\frac{\delta_{\text{ij}}R_{\text{z}}}{R^3} - \frac{\delta_{\text{iz}}R_{\text{j}}}{R^3} + \frac{\delta_{\text{jz}}R_{\text{i}}}{R^3} - \frac{3R_{\text{i}}R_{\text{j}}R_{\text{z}}}{R^5}\right),$$

$$\overline{\boldsymbol{y}} = \boldsymbol{y} - 2h\boldsymbol{e}_{\text{z}}, \qquad \boldsymbol{R} = R_{\text{i}} = \boldsymbol{x} - \overline{\boldsymbol{y}}, \qquad R = |\boldsymbol{R}|.$$

equation (4)

where $h$ is the nearest distance from the location of the force to a plane.

To prevent overlapping the swimmer and wall on the boundary surface, we assumed a linear repulsive force $\boldsymbol{F}^{\text{rep}}$ and a torque $\boldsymbol{T}^{\text{rep}}$ from the wall, defined as

$$\boldsymbol{F}^{\text{rep}} = \int_{\text{contact}} kl(\boldsymbol{r}^{\text{c}})\boldsymbol{e}_{\text{z}}\text{d}S(\boldsymbol{r}^{\text{c}})$$

$$\boldsymbol{T}^{\text{rep}} = \int_{\text{contact}} kl(\boldsymbol{r}^{\text{c}})\boldsymbol{e}_{\text{z}} \times (\boldsymbol{r}^{\text{c}} - \boldsymbol{r}'_{\text{g}})\text{d}S(\boldsymbol{r}^{\text{c}}),$$

equation (5)

where $l(\boldsymbol{r}^{\text{c}})$ is the immersed length between the location of node $\boldsymbol{r}^{\text{c}}$ belonging to a ciliary layer and the plane wall. A ciliary layer equivalent to the envelope of a ciliate was set at a distance $a$ from the boundary of the swimmer was at a distance $a$ from the boundary of the swimmer. The friction force between the wall and the swimmer was negligible. Here, assumption for the force and torque free conditions is expressed as the total force $\underline{\boldsymbol{F}^{\text{total}}}$ and the total torque $\boldsymbol{T}^{\text{total}}$ are zero. Using equation (1), the following equations are satisfied.

$$\boldsymbol{F}^{\text{total}} = \int_{\text{swimmer}} q_{\text{j}}^{\text{body}}(\boldsymbol{r}')\,\mathrm{d}S(\boldsymbol{r}') - \int_{\text{cilia}} F_{\text{j}}^{\text{cilia}}(\boldsymbol{r}'')\,\mathrm{d}S(\boldsymbol{r}'') - \boldsymbol{F}^{\text{rep}} = \boldsymbol{0}$$

$$\boldsymbol{T}^{\text{total}} = \int_{\text{swimmer}} q_{\text{j}}^{\text{body}}(\boldsymbol{r}') \times (\boldsymbol{r}' - \boldsymbol{r}'_{\text{g}})\,\mathrm{d}S(\boldsymbol{r}') - \int_{\text{cilia}} F_{\text{j}}^{\text{cilia}}(\boldsymbol{r}'') \times (\boldsymbol{r}'' - \boldsymbol{r}'_{\text{g}})\,\mathrm{d}S(\boldsymbol{r}'') - \boldsymbol{T}^{\text{rep}} = \boldsymbol{0}$$

equation (6)

Summarizing the equations (2), (3) and (6), we obtain the following relationships.

$$[\boldsymbol{u}_{\text{s}}] = [\boldsymbol{A}][\boldsymbol{q}^{\text{body}}]$$

$$[\boldsymbol{u}_{\text{s}}] = [\boldsymbol{B}][\boldsymbol{U}, \boldsymbol{\Omega}]$$

$$[\boldsymbol{F}^{\text{total}}, \boldsymbol{T}^{\text{total}}] = [\boldsymbol{C}][\boldsymbol{q}^{\text{body}}]$$

equation (7)

In this simulation, we discretized the swimmer surface into 5120 triangular meshes constructed from 2562 nodes, surface deformation was performed by displacing the nodes and imposing a nonslip boundary condition on the swimmer surface. After the discrimination of the surface integration with the triangles, the equation (7) becomes simultaneous equations. Then, accumulating the above three equations with a matrix,

$$\begin{bmatrix} \boldsymbol{0} \\ \boldsymbol{F}^{\text{total}}, \boldsymbol{T}^{\text{total}} \end{bmatrix} = \begin{bmatrix} \boldsymbol{A} & -\boldsymbol{B} \\ \boldsymbol{C} & \boldsymbol{0} \end{bmatrix} \begin{bmatrix} \boldsymbol{q}^{\text{body}} \\ \boldsymbol{U}, \boldsymbol{\Omega} \end{bmatrix}$$

equation (8)

The left term is known as force and torque free conditions $\boldsymbol{F}^{\text{total}} = \boldsymbol{0}$, $\boldsymbol{T}^{\text{total}} = \boldsymbol{0}$. In the right term, ***A, B*** and ***C*** are known. Therefore, by solving the inverse matrix with LU decomposition, the unknown parameters $\boldsymbol{q}^{\text{body}}$, ***U***, and ***Ω*** can be calculated. For the discrimination of the surface integration, the Gauss–Legendre quadrature and singularity integration methods were used to minimize the numerical error in the above integration (7, 8). A swimmer's location and orientation were updated from the solutions ***U*** and ***Ω*** by using a fourth-order Adams–Bashforth method at time step $\Delta t = 0.01$.

We set $\eta = 1$, $k = 2900$, $\underline{a} = 0.1$ and $|\boldsymbol{F}^{\text{cilia}}| = 161.28$. The distribution of $\boldsymbol{F}^{\text{cilia}}$ was like a band along the oral plane as shown in Fig. S8. The orientation of $\boldsymbol{F}^{\text{cilia}}$ was along the body surface as shown in Fig. S8. The location of $\boldsymbol{F}^{\text{cilia}}$ were located at a distance $2a/3$ from the boundary of the swimmer. Also, a flow filed in each time point is able to be calculated with the solution $\boldsymbol{q}^{\text{body}}$ in the equation (3). Our numerical swimmer model corresponded to a flow field qualitatively compared to a flow caused by *S. coeruleus* in terms of the number of vortices, vertex sizes, and locations.

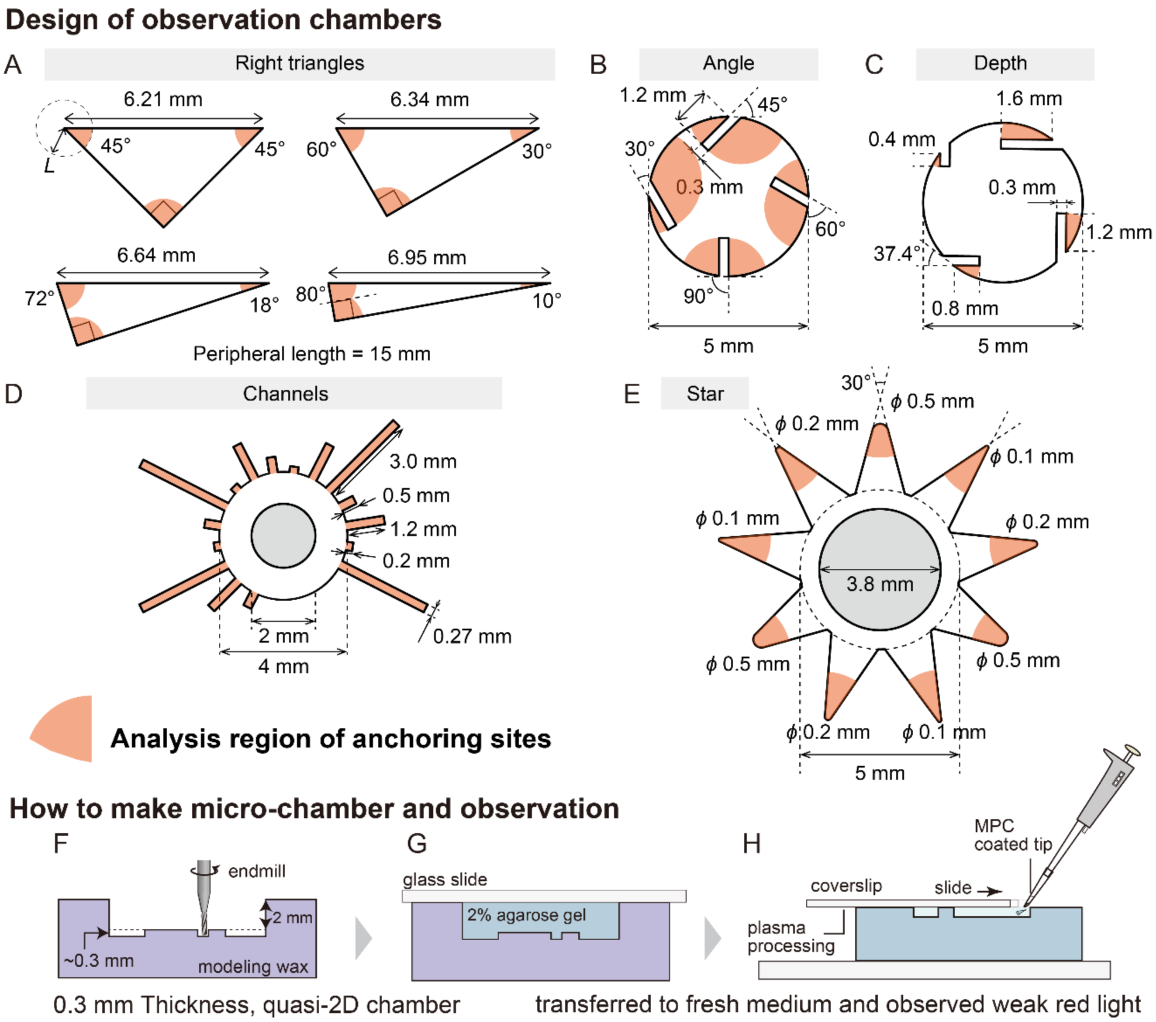


**Fig. S1. From design to observation in geometrical chambers.**

Top-view designs of the geometrical chambers containing the analyzed anchoring sites (A–E).

**(A)** Four types of right-triangle chambers with a 15 mm perimeter. The analysis regions were within 1 mm of each corner.

**(B)** Observation chamber with seven different corner angles formed by four protrusions from the outer edge. Angles less than 90° were measured between the protrusions and contact lines at the corner, and angles greater than 90° were estimated from supplementary angle for convenience.

**(C)** Observation chamber with four corners of varying depths formed by four protrusions from the outer edge. The protrusion length varied, but the angles were fixed.

**(D)** Four types of depth channels formed by 16 channels toward the outside from the outer edge.

**(E)** Three types of corners with different 2D curvatures but the same angle. The analysis regions of the anchoring sites were within *L'* (mm) = 2 + offset, which is the distance from the intersection of each corner.

**(F–H)** Side view schematics of fabrication and observation methods.

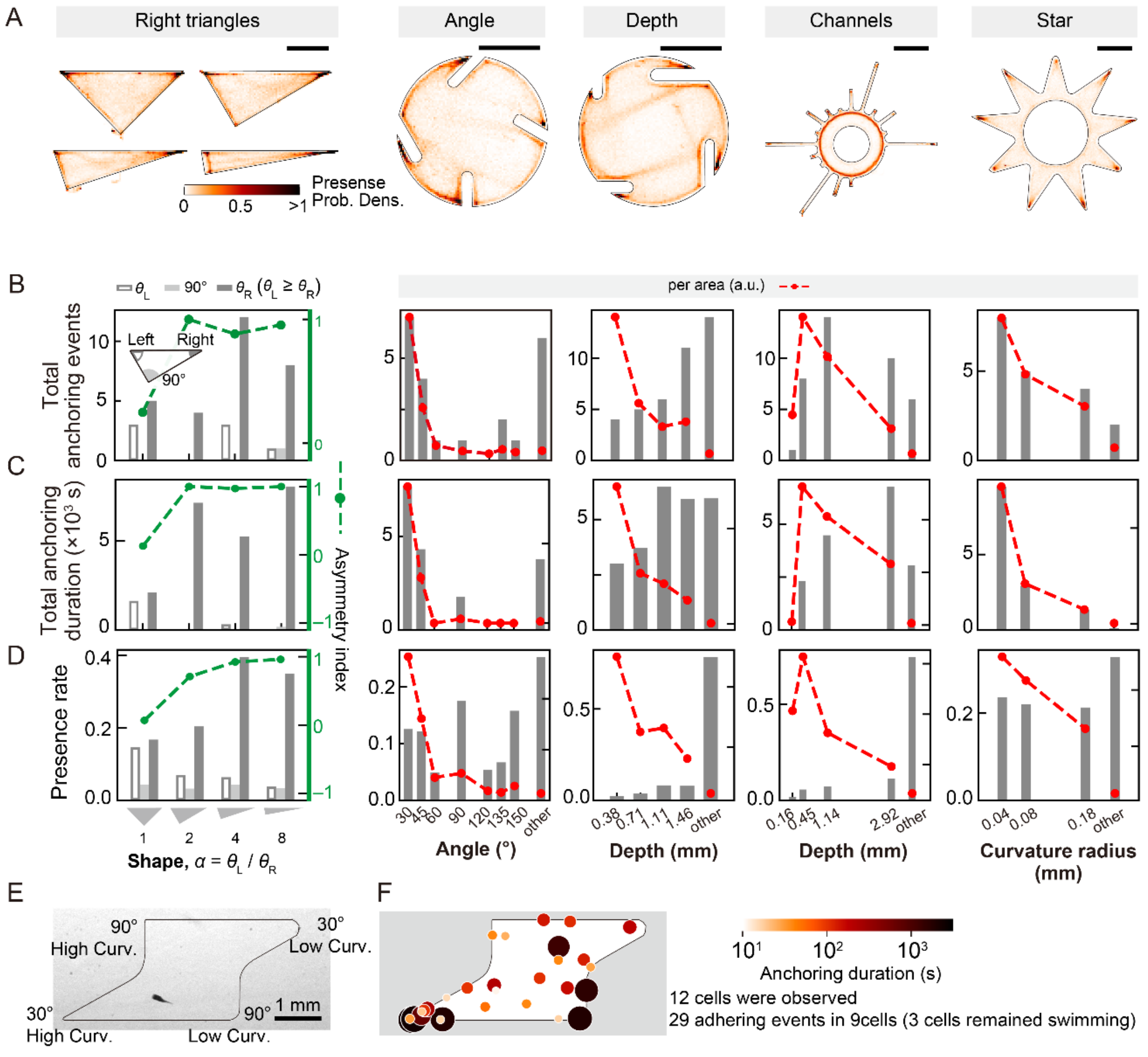


**Fig. S2. Behavioral analysis in the swimming state and anchoring duration in geometrical chambers.**

**(A)** 2D maps showing presence probability in the swimming state. Scale bars = 2 mm.
**(B)** The number of anchoring events in each geometrical region. The green and red dashed line represent the asymmetry index and each value per area (a.u.), respectively.
**(C)** Total anchoring duration at each geometrical region.
**(D)** Presence rates in each geometrical region, as defined in Fig. S1A–E.
**(E)** Top view of a geometrically structured chamber containing different corner angles and 2D curvatures.
**(F)** Anchoring sites and durations of 12 individual cells.

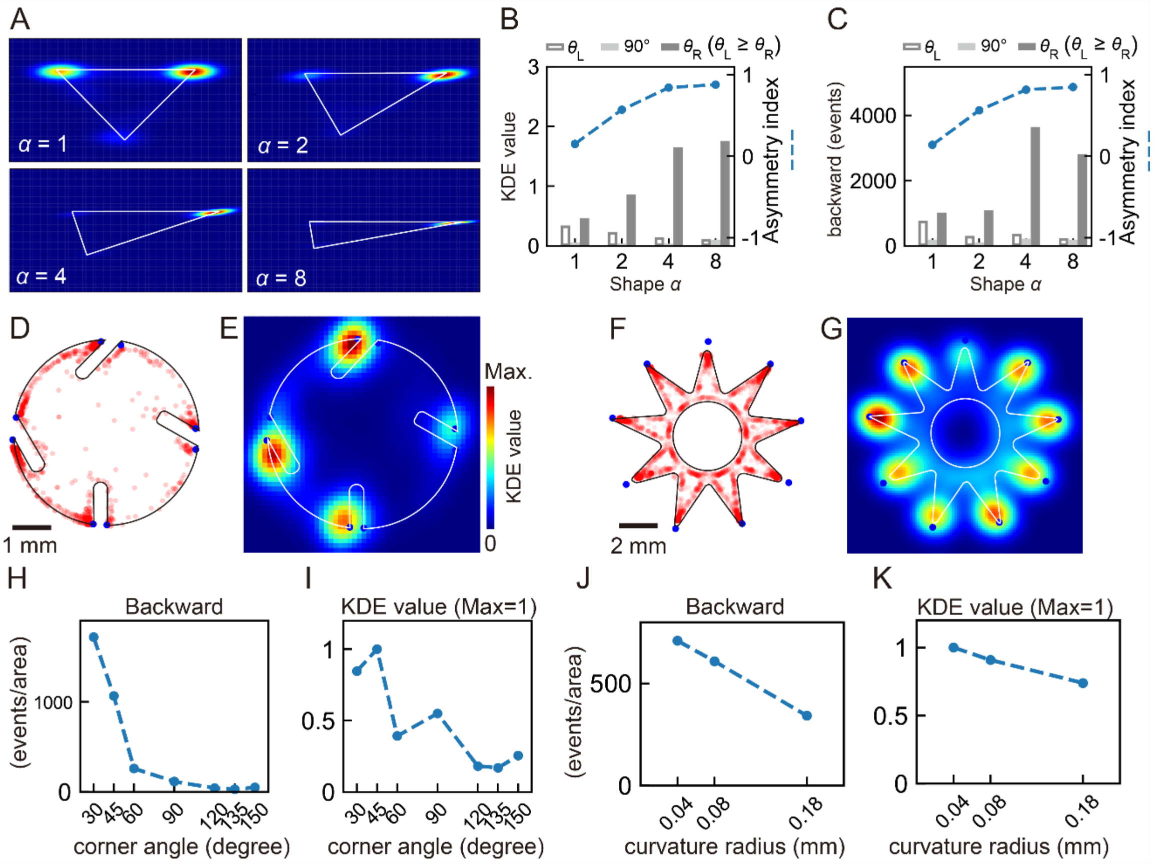


**Fig. S3. Backward swimming frequency related to geometrical features.**

**(A, E, G)** 2D maps of backward swimming frequency in geometrically structured chambers with different corner angles (A, E) and curvature (G). The maps were calculated using Kernel density estimation (KDE) of the backward swimming sites. White lines represent the outer edge of the chambers. To analyze the backward swimming event, we obtained the direction of the anterior end in the cell shape extracted from the cell contour and conducted binarization. From the asymmetric cell shape, we determined the anterior side. An inner dot between the anterior vector and the swimming vector was used to analyze the backward swimming.

**(B)** KDE values around the three corners relative to shape parameter *α*. The dashed line represents the asymmetry index.

**(C)** Number of backward events around the three corners relative to shape parameter *α*. The dashed line represents the asymmetry index.

**(D, F)** Locations of backward swimming are marked by red dots. The solid lines represent the outer walls of the chambers.

**(H, J)** Quantitative measurement of backward swimming per area.

**(I, K)** KDE values around each corner standardized to the maximum value in each chamber.

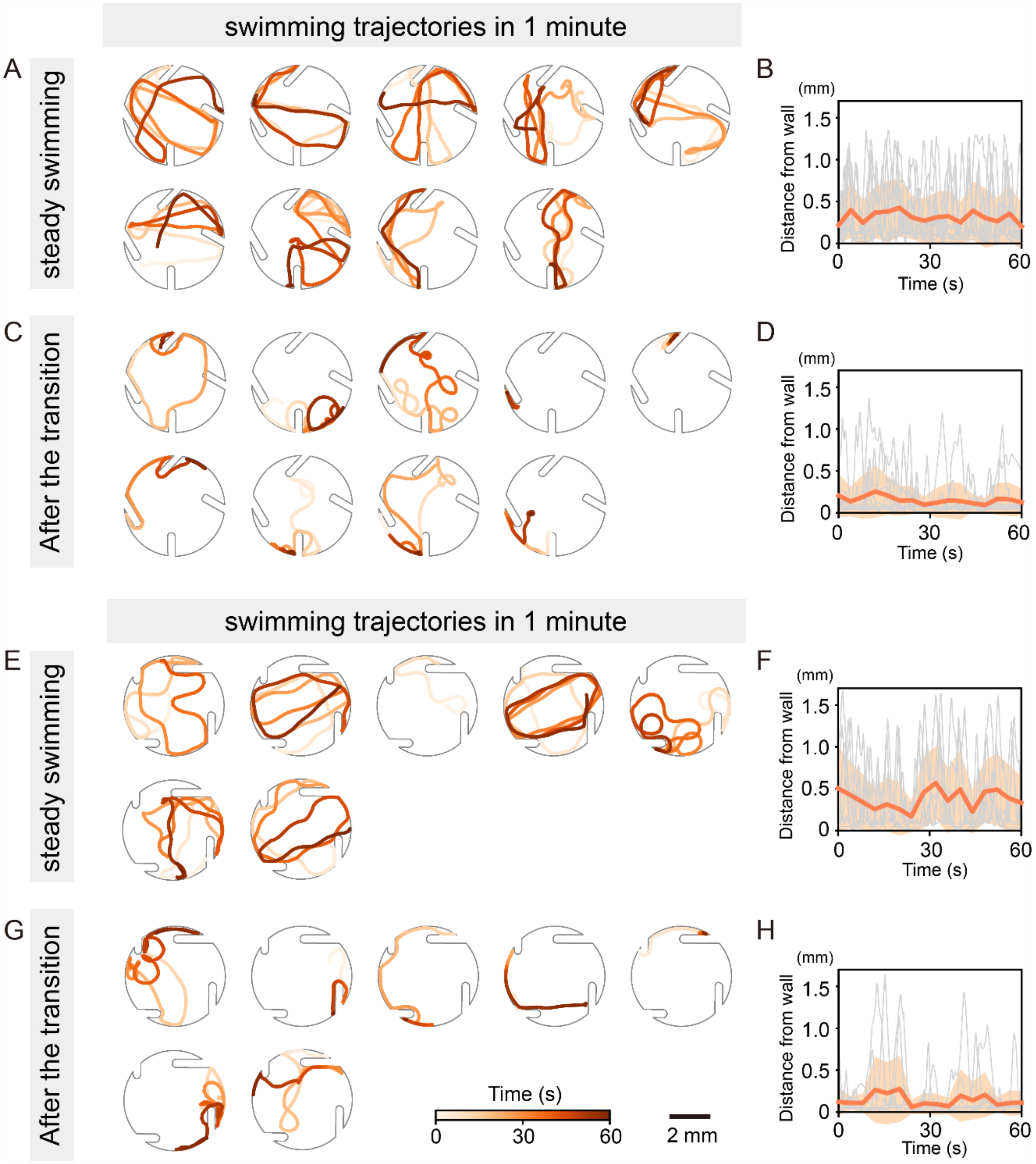


**Fig. S4. Swimming trajectories and the distance from the cell center to the wall before and after behavioral transitions.**

**(A, C, E, G)** Swimming trajectories in each chamber. Color represents time.

**(B, D, F, H)** Time series of the distance from the cell to the outer wall. Gray represents individual data. Solid orange lines and transparent orange bands denote the mean and SD, respectively.

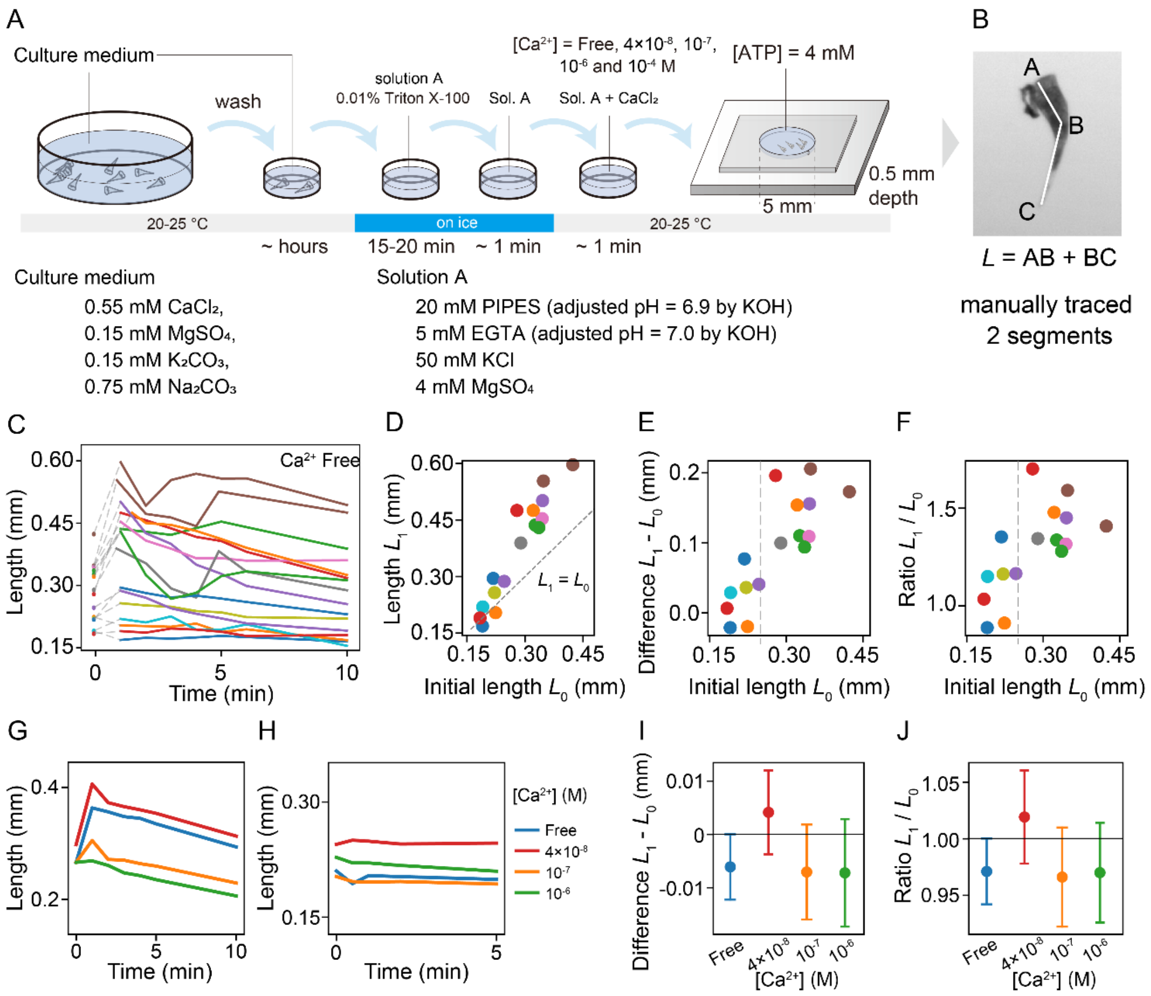


**Fig. S5. Effect of $Ca^{2+}$ concentration on cell deformation.**

**(A)** Schematic of detergent-treated cell model preparation.

**(B)** Model lengths measured from the total length of two manually traced segmented lines.

**(C)** Individual model length time series in $Ca^{2+}$-free conditions. ATP was added at time = 0.

**(D–F)** Model deformation in 1 minute and ATP addition based on the initial length $L_0$. Model lengths 1 min after ATP addition, $L_1$ (D), length differences (E), and deformation ratios (F) were compared with the initial length. The deformation in the below gray dashed line (initial length = 0.25 mm) was less than the model length above the gray dashed line.

**(G)** Time series of the mean model length at different $Ca^{2+}$ concentrations with ATP. ATP was added at time = 0.

**(H)** Time series of the mean model length at different $Ca^{2+}$ concentrations without ATP.

**(I, J)** Model deformation in 1 min without ATP. Model length differences (I) and deformation ratios (J) were compared with the $Ca^{2+}$ concentration. Deformation ratios were less than 5%. Dots and error bars represent the mean and SDs, respectively.

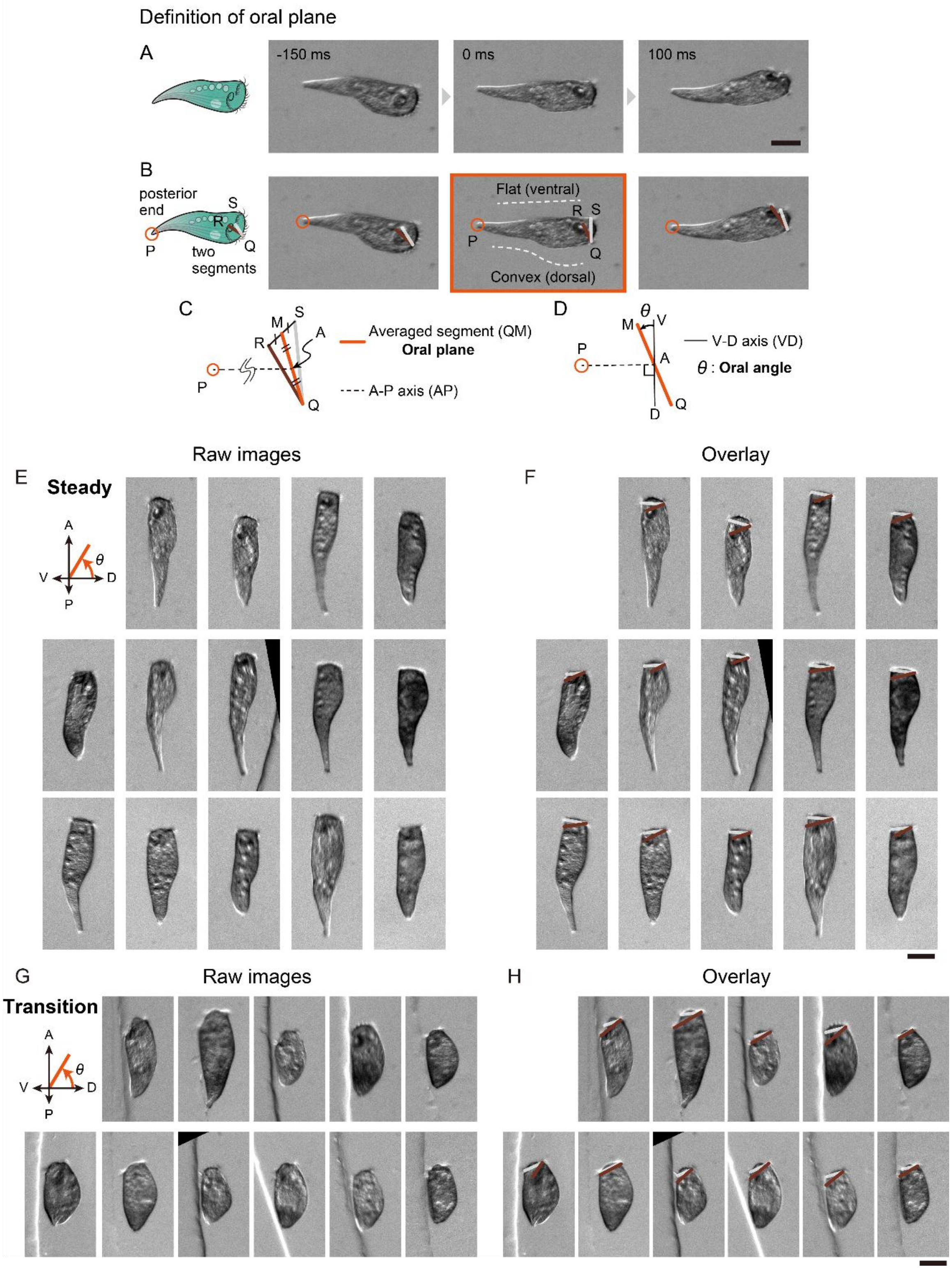


**Fig. S6. Measurement of the oral plane and individual images of the cell in the steady swimming state and after transition.**

**(A)** Schematic illustration of swimming *S. coeruleus* and sequential images at three time points.

**(B)** Manual detection of MB (brightness areas) forming the periphery of the three-dimensional oral plane. Two points from the ventral side (S and R, tips of MB) and one point from the dorsal side (Q) are shown. The image corresponding to the flat ventral side and convex dorsal side was used for measurement (image outlined by an orange solid rectangle).
**(C, D)** Geometry of the orientation angle of the oral plane. The oral angle was defined as the angle between the averaged segment QM, obtained from QS and QR, and the VD axis. The AP axis represents the segment between the posterior end (P) and the midpoint (A) of the averaged segment QM. The DV axis is orthogonal to the AP axis.
**(E–H)** Individual images of the cell in the steady swimming state (E) and after transition (G), aligned according to the AP and VD axes. The two segments QS and QR are overlaid in these images (F, H).
All scale bars are 0.1 mm.

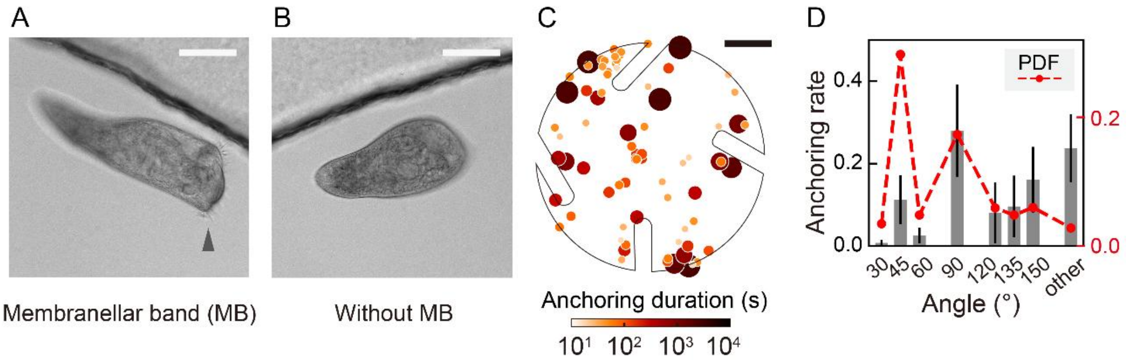


**Fig. S7. Anchoring sites of *S. coeruleus* without MB.**

Sucrose shock treatment was applied to intact *S. coeruleus* (A) with ciliary MB following Lin et al. (2018) (9). Treated *S. coeruleus* (B) shed MB around the anterior oral apparatus, resulting in a swimming speed approximately half that of intact cells. *S. coeruleus* without MB showed no geometrical preference in anchoring sites (13 cells across 4 experiments, mean ± SE) (C, D). In addition, wall-following behavior lasting 5 seconds was not observed for *S. coeruleus* without MB in the chamber. Scale bars are 0.1 mm for (A) and (B), and 1 mm for (C).

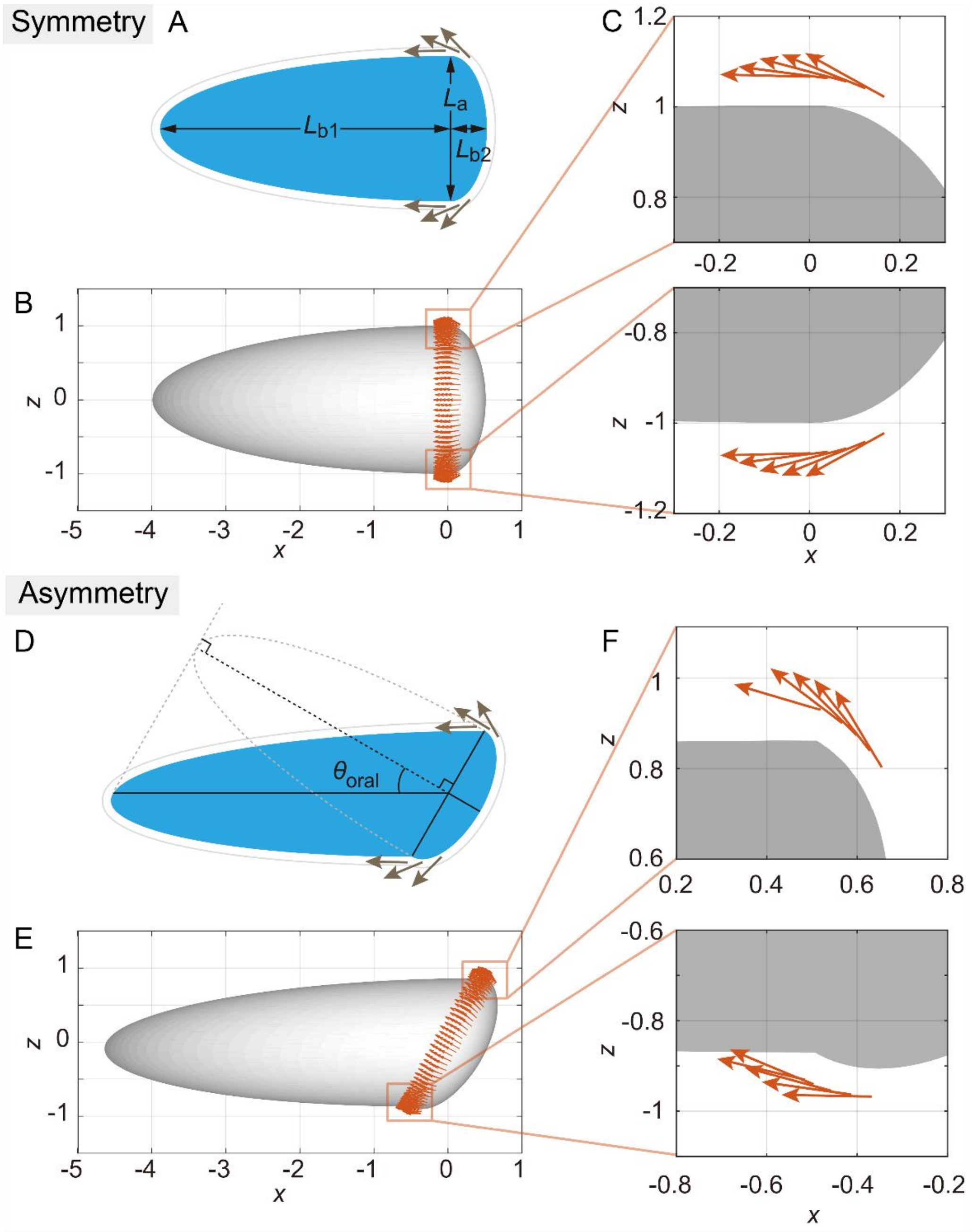


**Fig. S8. Deformation details and propulsive force distribution of two types of swimmers mimicking the *Stentor* cell.**

**(A)** A numerical swimmer was constructed from a sphere with radius $L_a/2 = 1$. One hemisphere was linearly elongated to $L_{b1} = 4$ times and the opposite hemisphere was compressed to half ($L_{b2} = 0.5$), respectively.

**(B, C)** The locations of point forces and closed view.

**(D)** Oral plane deformation was made by sliding the swimmer's posterior region, aligning the conical swimmer's base.

**(E, F)** The locations of the point forces with close-up views.

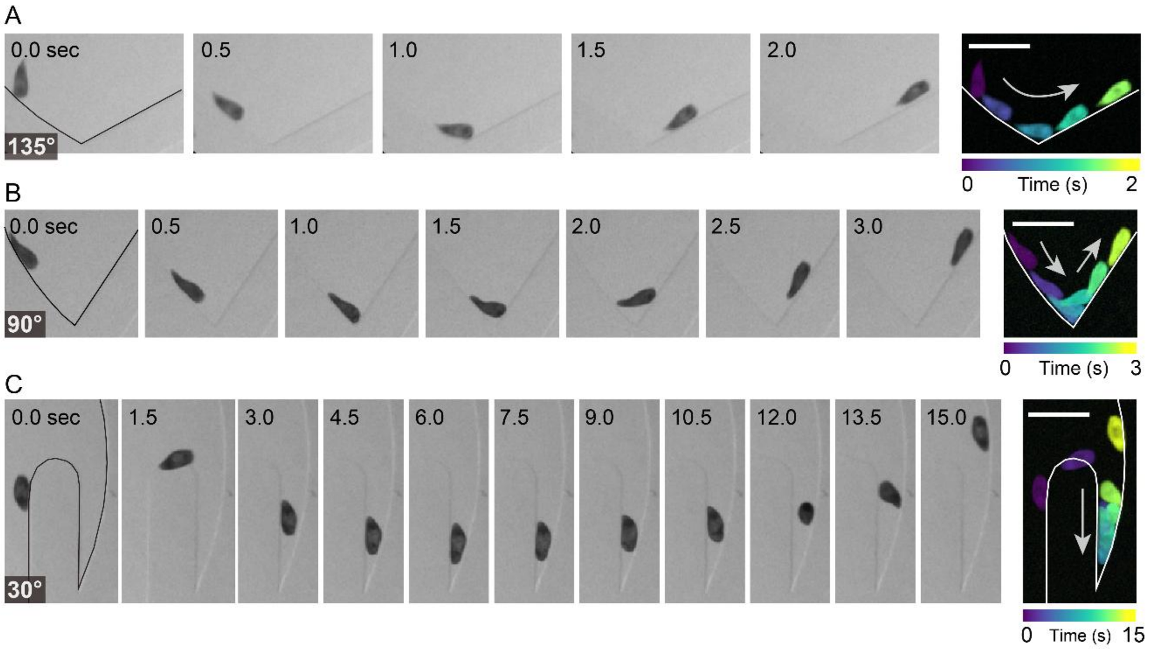


**Fig. S9. Contact interactions between wall-following *S. coeruleus* and the corner boundary across different angles, 135° (A), 90° (B), and 30° (C).**

Sequential images were acquired every 0.5 seconds in (A, B) or every 1.5 seconds in (C). A solid line was overlaid on each initial image to indicate the chamber boundary. Merged images were generated from the sequential images. Color represents time. All scale bars are 0.5 mm. See also Movies S5–S7.

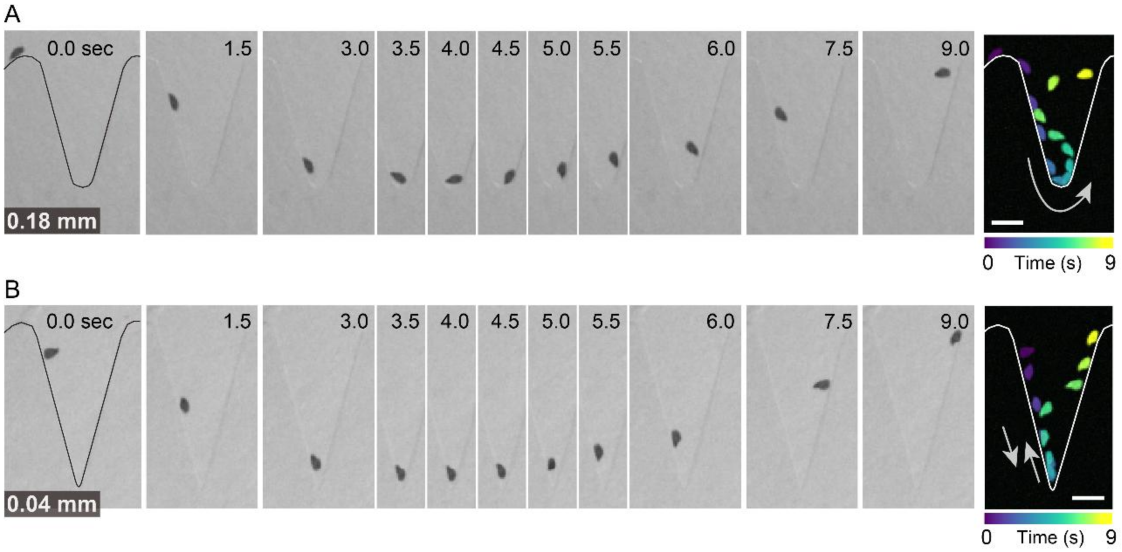


**Fig. S10. Contact interactions between wall-following *S. coeruleus* and the corner boundary across different curvature radii, 0.18 mm (A) and 0.04 mm (B).**

Time is represented by the labels above each image in the sequential frames. A solid line was overlaid on each initial image to indicate the chamber boundary. Merged images were generated from sequential images acquired at 0.75-second intervals. Color represents time. All scale bars are 0.5 mm. See also Movies S8 and S9.

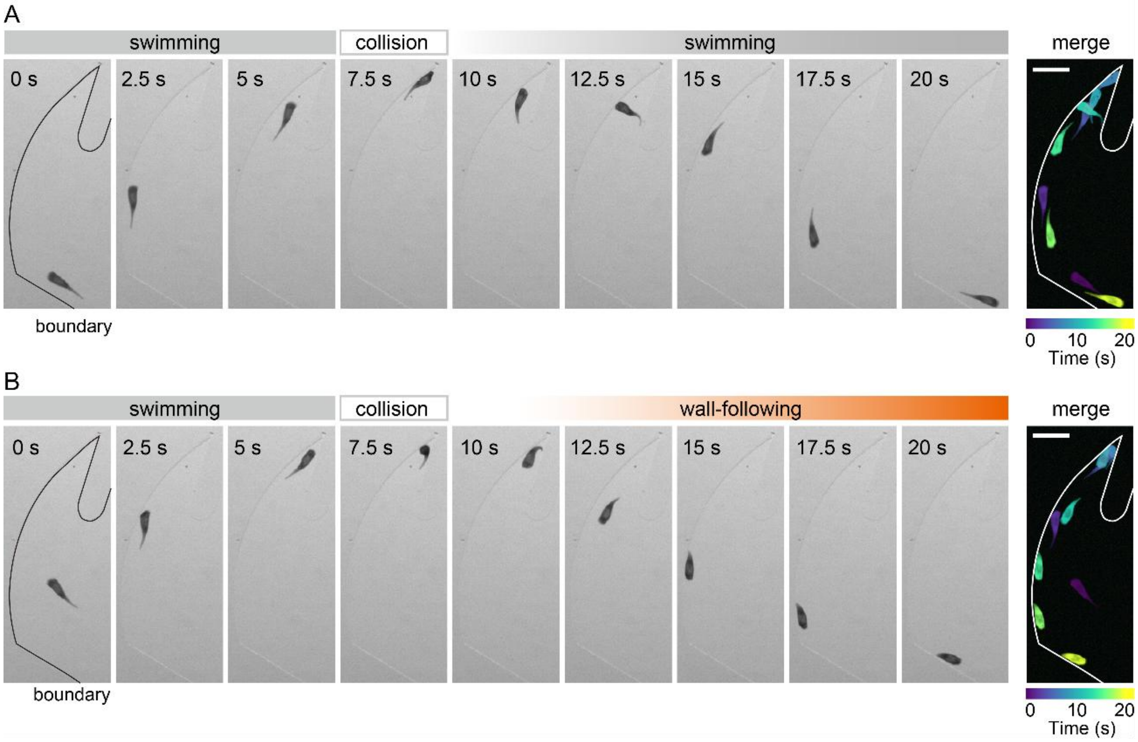


**Fig. S11. Two collision processes in a geometrically structured chamber.**

Sequential images were acquired every 2.5 seconds. A solid line was overlaid on each initial image to indicate the chamber boundary. Merged images were generated from the sequential images. Owing to the confined observation chamber, collisions were frequently observed during the steady swimming state, and the cell remained in that state (A). A few seconds before the onset of wall-following behavior, the cell collided with the corner wall (B). Color represents time. All scale bars are 0.5 mm. See also Movies S10 and S11.

#### Legends for Movies S1 to S11

**Movie S1. Anchoring behavior in *S. coeruleus*, related to Fig. 2.**
A swimming *S. coeruleus* swims in the structural geometrical chamber. After the cell changes its motility, the cell finally anchors in a narrow space. A solid line was overlaid on the movie to indicate the chamber boundary. Played at 5× speed. Scale bar = 1 mm.

**Movie S2. Typical swimming behavior of *S. coeruleus* in the steady swimming state, related to Fig. 3.**
A conical swimming *S. coeruleus* in the geometrically structured chamber. The straightforward movement with a spiral trajectory is influenced by the boundary of the chamber's geometry, and the cell explores the whole region. The swimming trajectory in Fig. 3C was obtained from this movie. A solid line was overlaid on the movie to indicate the chamber boundary. Playback at 4× speed. Scale bar = 1 mm.

**Movie S3. Typical wall-following behavior in the transition before anchoring in *S. coeruleus*, related to Fig. 3.**
A swimming *S. coeruleus* with a tilted oral apparatus moves along the wall of the observation chambers. The swimming trajectory in Fig. 3C was obtained from this movie. A solid line was overlaid on the movie to indicate the chamber boundary. Playback at 4× speed. Scale bar = 1 mm.

**Movie S4. Motions of the numerical *Stentor* near a nonslip surface, related to Fig. 5.**
The symmetric conical swimmer ($\theta_{\text{oral}}$ = 0°) reflects off (top), and the asymmetric conical swimmer ($\theta_{\text{oral}}$ = 30°) moves along (bottom) the nonslip surface located at $z$ = 0. The center of the swimmer is initially located at ($x$, $y$, $z$) = (0, 0, 2), and the anterior plane is oriented to the surface at ~30°.

**Movie S5. Contact interaction between wall-following *S. coeruleus* and a 135-degree corner, related to Fig. S9A.**
Wall-following *S. coeruleus* mechanically interacts with a 135-degree corner. Playback at 2× speed. Scale bar = 0.5 mm.

**Movie S6. Contact interaction between wall-following *S. coeruleus* and a 90-degree corner, related to Fig. S9B.**
Wall-following *S. coeruleus* mechanically interacts with a 90-degree corner. Playback at 2× speed. Scale bar = 0.5 mm.

**Movie S7. Contact interaction between wall-following *S. coeruleus* and a 30-degree corner, related to Fig. S9C.**
Wall-following *S. coeruleus* mechanically interacts with a 30-degree corner. Playback at 2× speed. Scale bar = 0.5 mm.

**Movie S8. Contact interaction between wall-following *S. coeruleus* and a corner with a curvature radius of 0.18 mm, related to Fig. S10A.**
Wall-following *S. coeruleus* mechanically interacts with a corner with a curvature radius of 0.18 mm. Playback at 2× speed. Scale bar = 0.5 mm.

**Movie S9. Contact interaction between wall-following *S. coeruleus* and a corner with a curvature radius of 0.04 mm, related to Fig. S10B.**

Wall-following *S. coeruleus* mechanically interacts with a corner with a curvature radius of 0.04 mm. Playback at 2× speed. Scale bar = 0.5 mm.

**Movie S10. Collision at a 30-degree corner during steady swimming, related to Fig. S11A.**

*S. coeruleus* collided with a 30-degree corner in a steady swimming state. The cell then turned and reversed direction, swimming steadily again in the opposite direction. Playback at 2× speed. Scale bar = 0.5 mm.

**Movie S11. Transition from steady swimming to wall-following immediately before corner collision, related to Fig. S11B.**

*S. coeruleus* collided with a 30-degree corner in a steady swimming state. The cell then contracted and followed the chamber boundary. Playback at 2× speed. Scale bar = 0.5 mm.
